# Genome-Wide in silico analysis reveals activation of a silent resistome driving imipenem resistance in *Pseudomonas aeruginosa*

**DOI:** 10.64898/2026.01.25.701575

**Authors:** Sarfraz Anwar, A R Aromal, Ananya Anurag Anand, Sintu Kumar Samanta

## Abstract

Resistance to imipenem in *Pseudomonas aeruginosa* relies on multiple factors that remain poorly understood. In our work, we performed a systemic analysis of genome-wide changes involved in resistance in a set of 95 clinically unrelated strains, including 41 resistant (MIC ≥ 64 mg/L) and 54 susceptible (MIC ≤ 2 mg/L) isolates. Our approach is based on the pan-genomics analysis, combining the use of core-genome phylogenetic analysis, MLST (Multilocus Sequence Typing), GWAS (Genome-Wide Association Studies) and variant level profiling of the blaOXA genes. Higher-order structure within the set was studied using methods of the co-occurrence networks and WGCNA (weighted gene co-expression network analysis) specifically adjusted to handle presence/absence data. Despite having a broader and more diverse resistome, no clonal grouping of the resistant isolates was observed indicating independent evolutionary origins. The LASSO model using a lineage-aware approach showed robust predictive capability (AUC = 0.836) that validates the polygenic characteristic of resistance. Twelve accessory genes were found to be significant determinants of resistance; however, only four genes (group_10880, group_10887, group_4947, and phzB) were identified using both GWAS and gene network analysis, showing involvement in protein folding, metal stress response, genome plasticity, and metabolic adaptation. Interestingly, some carbapenemase-active variants of blaOXA were also found in imipenem-susceptible strains, showing that gene presence alone does not ensure resistance. We therefore propose the “Silent Resistome Activation Model,” where resistance genes become functional only with support from identified accessory genes and coordinated interactions at both the genomic and network levels.

## 1. Introduction

*Pseudomonas aeruginosa* is a significant public health problem worldwide, with 559,000 deaths annually, of which more than half are due to antimicrobial resistance (AMR) **(Al-Daghistani et al., 2025).** Antimicrobial resistance is a major public health issue that has also come to be recognized as a critical threat, causing 700,000 deaths worldwide annually and expected to escalate to 10 million deaths annually by 2050, in addition to the economic impact. The United States also reported 32,600 hospital-associated *P. aeruginosa* infections and 2,700 deaths due to the same in 2017, as per surveillance data provided by the Centers for Disease Control and Prevention (CDC). A significant increase in the incidence of *P. aeruginosa* has also been reported in the subsequent years. *P. aeruginosa*, in light of its public health importance, was included in the list of the seven most important antimicrobial-resistant pathogens commonly encountered in the healthcare setting in 2024 by the CDC.

Recently, a meta-analysis study indicated that the global burden of carbapenem-resistant *P. aeruginosa* (CRPA) is high, with the pooled prevalence of CRPA among clinical isolates being approximately 34.7% **(Ramatla et al., 2025).** The study also indicated that the prevalence of CRPA is high in some regions of Europe and South America, while studies from Asia, Africa, and North America indicated the significant presence of CRPA, which suggests the global distribution of the pathogen in various geographical locations.

Imipenem, a broad-spectrum carbapenem, still holds a central place in the empirical treatment of severe and life-threatening infections caused by the bacterium *P. aeruginosa* **(Onguru et al., 2008).** Nevertheless, a rise in resistance to imipenem has been noted in various regions of the world, as classified by the World Health Organization (WHO), where resistance rates have surpassed 30% **(Ahmadi et al., 2025).** Nonetheless, the critical role imipenem plays in the clinic does not support the fact that imipenem non-susceptibility in *P. aeruginosa* is linked to a specific and consistent genetic marker (**Shields et al., 2022**). Instead, the mechanisms that underlie imipenem resistance in *P. aeruginosa* are complex and multifactorial, including the overexpression of efflux pumps, the absence or alteration of outer membrane porins, the acquisition of carbapenemases, and biofilm-related tolerance. However, the contribution of each mechanism to imipenem resistance in *P. aeruginosa* is a topic of debate **(Alvarez-Ortega et al., 2010; Musafer et al., 2014; Simanek et al., 2022).**

The majority of previous genomic studies on *P. aeruginosa* have been restricted to a small number of reference genomes or particular outbreak-associated strains. These studies have given us important insights, but it is hard to distinguish between resistance characteristics that can be considered common for the species (core genome) and those that differ between strains (accessory genome) **(Pajaro-Castro et al., 2025; Gómez-Martínez et al., 2023; Kung et al., 2010).** Therefore, only a few studies have used pangenome-scale approaches for large numbers of well-characterized susceptible and resistant strains, especially with the aim of determining their relationship with virulence-associated characteristics (**López-Pérez et al., 2021**). Moreover, genome-wide association studies (GWAS) comparing imipenem-susceptible and imipenem-resistant *P. aeruginosa* isolates remain limited. To date, there has also been little investigation at the clonal level to determine which sequence types (STs) are preferentially associated with imipenem resistance in *P. aeruginosa*. Importantly, the integration of machine learning–based prediction and network-level approaches such as WGCNA with pangenome data has rarely been explored.

To address these gaps, 95 epidemiologically independent *P. aeruginosa* strains were investigated using an integrated genomic approach, with resistance to imipenem (≥64 mg/L, n = 41) and susceptibility to imipenem (≤2 mg/L, n = 54) strains obtained from the Pathosystems Resource Integration Center (PATRIC) repository. Phenotypic validation confirmed accurate categorization of the strains. Pangenome analysis, core-genome phylogenies, MLST, and pan-genome wide-association study with phylogenetic adjustment, as well as machine learning with leave-one-sequence-type-out validation for lineage-specific prediction using LASSO algorithm, have been employed. Gene co-expression networks and WGCNA helped to identify several modules associated with resistance, involving protein folding, metal stress responses, genome plasticity, and metabolic adaptation. Interestingly, the presence of various types of blaOXA genes, the main genes associated with carbapenemase activity, was found among the susceptible isolates, confirming that not only presence of the genes is responsible for resistance, but also the genomic environment around those genes.

## 2. Materials and Methods

### 2.1 Sample Selection

After applying several filtering steps, we identified 95 high-quality genomes for our analysis. We began with all available *P. aeruginosa* samples in the PATRIC database **(Davis et al., 2020)** and then applied strict criteria to separate imipenem-resistant and imipenem-susceptible isolates. For the resistant group, we kept only those isolates with an MIC value ≥ 64 mg/L, while for the susceptible group, we selected isolates with an MIC value ≤ 2 mg/L. For ensuring that all samples were laboratory-validated, it was required that MIC was determined using either agar dilution or broth dilution methods **(Wiegand et al., 2008)**. Along with the quality filter based on MIC, several other quality filters were implemented, such as removing low-quality assemblies, duplicates, and samples with poor CheckM quality and assembly statistics **(Parks et al., 2015).** Finally, the dataset was comprised of 41 resistant and 54 sensitive genomes, making up a total of 95 samples. These samples were either complete genome assemblies or high-quality WGS.

### 2.2 Identifying Antibiotic Resistance Genes (ARGs)

To identify the genes responsible for antimicrobial resistance within the *P. aeruginosa* genome, AMRFinderPlus version 3.11.17 was used **(Feldgarden et al., 2021)**. This program was designed and maintained by the National Center for Biotechnology Information (NCBI). AMRFinderPlus works by using the reference gene database, which is curated and of high quality, using Hidden Markov models.

### 2.3 Identifying Virulence Factors (VFs)

To identify virulence-associated genes within the *P. aeruginosa* genomes, ABRicate (version 1.0.1) was used (https://github.com/tseemann/abricate), which is a commonly used tool for genome screening. For this study, the virulence factor database (VFDB), which is often used in bacterial pathogenicity studies, was chosen as a reference database, as described by **Chen et al. (2016)**. ABRicate scans genomes by comparing each genome’s sequence to a reference database, VFDB, and identifying genes associated with known virulence factors. This allowed for the identification of virulence factors in all genomes included in this study.

### 2.4 Pan-genome construction and phylogenetic analysis

Pan-genome analyses were performed separately for imipenem-resistant and imipenem-susceptible *P. aeruginosa* isolates, and subsequently for the complete set of 95 genomes. Prior to pan-genome construction, all genome assemblies were re-annotated using Bakta (version 1.9.1) **(Schwengers et al., 2021).** As Bakta employs an alignment-free annotation approach supported by its own local reference database, the full database (∼31.9 GB) was downloaded to ensure consistent and high-quality gene annotations across all genomes.

The re-annotated genomes were analysed using Panaroo (version 1.5.2) with default parameters **(Tonkin-Hill et al., 2020).** Panaroo applies a graph-based framework that corrects common annotation errors and effectively resolves paralogous gene clusters. The standard workflow included strict cleaning, gene clustering using CD-HIT at 98% sequence identity and merging of gene families based on neighbourhood context at a 70% identity threshold **(Tonkin-Hill et al., 2020).**

From the full dataset, a core genome alignment was produced for all 95 isolates using the program MAFFT, version 7.526, which is the integrated multiple sequence alignment tool within the Panaroo **(Katoh et al., 2019).** A phylogenetic tree was produced using the concatenated core gene alignment, and this was done using the program IQ-TREE, version 3.0.1 **(Nguyen et al., 2015).** The best substitution model was determined using the Bayesian Information Criterion, and ultrafast bootstrap analysis was used for assessing node support, using 1,000 bootstrap replicates **(Minh et al., 2013; Trifinopoulos et al., 2016)**. The phylogenetic tree was visualized and annotated using the iTOL web server **(Letunic & Bork, 2024).**

### 2.5 Pan-GWAS analysis

To determine genetic factors linked to imipenem resistance, we performed pangenome-wide association analysis using two approaches. Scoary was used to control for phylogenetic relationships between the isolates, as described by **Roder and colleagues**, and Pyseer was used to apply a linear mixed model, which reduces the impact of population structure, as described by **Lees and colleagues (Less et al.,2018; Roder et al., 2024).** Using both methods enabled the robust identification of accessory genes that may be linked to resistance.

For both tests, imipenem’s susceptibility status, either resistant or susceptible, was set as the response variable, and the gene presence/absence matrix, which was generated by Panaroo, was set as the input dataset. Scoary was run with default parameters and tested the distribution of each gene between resistant and susceptible populations. Genes with a Benjamini-Hochberg-adjusted p-value of less than 0.05 and an odds ratio of more than 1 were selected as putative genes associated with resistance (Bhalla & Nanda, 2024; Ortega-Sanz et al., 2025).

The analysis was also performed using Pyseer, version 1.3.1. To account for population structure, a core-genome distance matrix was included as a kinship matrix, and significant associations were determined by performing a likelihood ratio test with an adjusted p-value cutoff set at 0.05 (Redfern et al., 2021). For each gene that was found to be significant, the associated beta coefficient was also noted, which would indicate the strength and direction of association with imipenem resistance.

### 2.6 Identification of MLST Profiles and ST-Level Comparison of ARGs and VFs

To analyze the variability of the antibiotic resistance genes (ARGs) and virulence factors (VFs) in different genetic lineages, the *P. aeruginosa* isolates, which were resistant and susceptible to imipenem, were subjected to the multilocus sequence typing (MLST) technique. The sequence type (ST) for each isolate was determined by the MLST command line tool version 2.23.0, which identifies the ST based on the seven core genes, known as the housekeeping genes **(Page et al., 2017).** After the ST assignment, the isolates were grouped based on their sequence type, and the composition of ARGs and VFs was compared. The average number of resistance genes and virulence genes per ST was calculated, which helped to identify the sequence types that were enriched with either of the two gene categories. As an illustration of these patterns, a comparative plot was developed based on the mean abundance of ARGs and the mean abundance of VFs for each sequence type. This exercise was useful for identifying STs that potentially contributed more to imipenem resistance due to the presence of resistance and virulence-associated genes.

### 2.7 Machine Learning-Based Resistance Prediction

For the construction of a predictive model for imipenem resistance in *P. aeruginosa*, the technique of LASSO logistic regression, based on the gene presence/absence data, was used **(Tibshirani, 1996).** The model was specifically designed to determine if a small number of genetic characteristics could accurately differentiate between resistant and susceptible isolates.

#### 2.7.1 Data Preparation and Feature Engineering

Input data consisted of a binary gene presence/absence data set, as well as strain metadata such as sample identifiers, resistance phenotypes, and sequence type information. Candidate genes were chosen based on a set of twelve genes that had been previously used as predictive features from genome-wide association screening data. The data was represented as a set of binary values for the presence or absence of the genes in each strain, with the target variable represented as a set of binary values for resistance status, 1 for resistant strains and 0 for susceptible strains **(Sunuwar & Azad, 2022).**

Sequence type information was also handled with care in order to avoid any leakage of information during cross-validation. Unknown or missing ST assignments, which were represented in different forms as “unknown,” were replaced with unique identifiers “UNK_01,” “UNK_02,” etc. (Wilimitis & Walsh, 2023).

All features were standardized using z-score standardization, where all features are scaled to have zero mean and unit variance, making it easier for the models to understand and compare the scales of the different variables (Pinheiro et al., 2025; Wilimitis & Walsh, 2023).

#### 2.7.2 LASSO Logistic Regression and Hyperparameter Tuning

We defined the L1 regularized logistic regression model with a number of important parameters. The solver used was ‘liblinear’, which is efficient in its computation when the dataset is small to medium in size and also supports L1 regularization **(Fan et al., 2008).** Class weights were defined as ‘balanced’, which would automatically compensate if there were more resistant than susceptible isolates in the data and prevent the model from becoming imbalanced and predicting the most prevalent class **(He & Garcia, 2009; King & Zeng, 2003).**

LogisticRegressionCV class with 5-fold cross-validation was used on twenty possible values of C on a logarithmic scale (Varma & Simon, 2006; Mahlich et al., 2024). The five-fold cross-validation scheme is commonly used in genomic prediction problems and is stable in selecting the hyperparameters without incurring unnecessary computational costs (Alemu et al., 2024). This process selected the optimal value of C that maximized the performance while ensuring the model’s parsimony by incorporating L1 shrinkage.

#### 2.7.3 Model Training and LOSO (leave-one-sequence-type-out) Cross-Validation

An important factor to consider during bacterial genomic prediction is the population structure. During k-fold cross-validation, the data is split into the training and test sets, which does not guarantee that the two sets do not have similar data points (i.e., the same sequence type). This is not a realistic approach, as the model is actually being taught the specific characteristics of the data rather than the actual mechanisms of the data **(Yu et al., 2025).**

Therefore, we adopted leave one sequence type out cross-validation as our primary validation method (Cheng et al., 2017). This involves the exclusion of all isolates of a given sequence type as a validation set, and the model is trained on all remaining sequence types. This is repeated iteratively until each sequence type has been used as a validation set once. This method ensures that the model is validated on sequence types that are genetically distinct and were not included in the model during the training phase.

For each of the held-out STs, we fit a new LASSO model using the previously found optimal value of C, made predictions for the probability of resistance, and then used a threshold of 0.5 to create binary predictions (Orcales et al., 2024; Wiesmueller et al., 2026). We recorded the true labels, predicted probabilities, and binary predictions for all isolates across all folds.

#### 2.7.4 Performance Evaluation

Quantification of model discrimination ability was performed by measuring the area under the receiver operating characteristic curve, abbreviated as AUC-ROC **(Nahm, 2022; Çorbacıo**ğ**lu & Aksel, 2023).** Additional metrics used to characterize the performance of the classification were the sensitivity (true positive rate), specificity (true negative rate), positive predictive values, negative predictive values, and overall accuracy **(Hicks et al., 2022).** These metrics were also computed using the aggregated confusion matrix over all LOSO folds. We also report the per-ST performance where there were sufficient samples and outcome variation within each sequence type to calculate meaningful AUC values.

#### 2.7.5 Feature Importance and Interpretation

LASSO is a type of Regularization that performs both Feature Selection and Shrinkage at the same time **(Tibshirani, 1996).** It sets the coefficients of less important features closer to zero.

We used the final coefficients of each machine learning model as a method of determining the genes that are most associated with the prediction of resistance **(Akter et al., 2025).** This is because the absolute value of each coefficient represents the strength of association after adjusting for all other features.

All machine learning computations were done using scikit-learn **(Pedregosa et al., 2012).**

### 2.8 Univariate Gene-Level Diagnostic Evaluation

To assess the individual contribution of each gene to the imipenem resistance, gene-wise diagnostic analysis has been conducted **(Xu & Setiono, 2015).** In this approach, the individual genes are evaluated based on their absence/presence status, and the clear picture of the genes with strong individual association with the resistance phenotypes is obtained.

#### 2.8.1 Data Integration and Preparation

We also imported strain metadata and gene presence-absence matrices, where the sample identifiers were standardized as strings **(Choudhury & Andam, 2026).** The merged data included the resistance status as the standard and the binary gene presence indicators as the predictive features. All genes in the matrix were included in the assessment, regardless of their known association status.

#### 2.8.2 Confusion Matrix Construction

For each gene, a 2 x 2 matrix was constructed crossing over the predicted resistance status based on the presence of the gene and the observed resistance phenotype. The elements of the 2 x 2 matrix were defined as TP (resistant strains with the gene), FP (susceptible strains with the gene), TN (susceptible strains without the gene), and FN (resistant strains without the gene) **(Moradigaravand et al., 2018).** Strains from which the gene was either completely absent or present in all strains were excluded from the calculation as these did not provide any discriminatory information.

#### 2.8.3 Performance Metrics Calculation

For each of the confusion matrices, three diagnostic performance metrics are derived. Precision or the positive predictive value was derived as TP/(TP+FP), the probability that a strain carrying the gene is indeed resistant **(Condorelli et al., 2024).** The sensitivity or true positive rate was derived as TP/(TP+FN), the proportion of resistant strains correctly identified as such by the presence of the gene. The specificity or true negative rate was derived as TN/(TN+FP), the proportion of susceptible strains correctly identified as such by the absence of the gene.

#### 2.8.4 Gene Ranking and Prioritization

Ranking of the genes was performed based on the number of false positives, ranking those genes with the least number of false positives as the first priority, i.e., those that are least likely to misclassify susceptible strains as resistant **(Ali et al., 2018; Mukiri et al., 2025).** In cases where the number of false positives was the same for more than one gene, precision values were used as a secondary criterion for ranking in descending order **(Kim et al., 2022).** This ranking strategy reflects clinical priorities where minimizing false alarms (unnecessary concern for susceptible isolates) may outweigh maximizing sensitivity, particularly in screening contexts.

### 2.9 Gene Co-occurrence Network Construction and Hub Gene Identification

A gene co-occurrence network was generated from the pan-genome gene presence–absence matrix together with strain phenotype metadata (resistant/susceptible). First, the matrix was aligned with the metadata and genes with very low prevalence were removed by keeping only those present in at least a minimum number of strains in either phenotype group. The filtered dataset was then divided into resistant and susceptible subsets. Pairwise Spearman correlation coefficients were calculated between all gene pairs separately for each group using a vectorized approach, and p-values were computed and corrected for multiple testing using the Benjamini– Hochberg false discovery rate (FDR) method **(Benjamini & Hochberg, 1995, Gautreau et al., 2020; Spearman, 1904).** The resulting correlation and adjusted p-value matrices were exported as input files for downstream WGCNA module analysis. Network edges were defined using an absolute correlation cutoff of ≥ 0.3 and FDR-adjusted p-value < 0.05 **(Johansson et al., 2011; Randhawa & Acharya, 2015).** A resistant-specific network was then constructed by selecting only those gene pairs that were significant in resistant strains and either not significant in susceptible strains or showed at least 0.2 higher absolute correlation in the resistant group **(Akoglu, 2018).** This resistant-only edge list was used to build the final network. The node connectivity, also known as the degree, of the network was calculated, and the genes were ranked based on their degree centrality. The top 10 most connected genes were selected as the hub genes, and a simplified edge file for the hub-only network was generated for later visualization and interpretation **(Song et al., 2020).**

### 2.10 Weighted Gene Co-expression Network Analysis (WGCNA)

#### 2.10.1 Data Preprocessing

Weighted gene co-expression network analysis was performed to find the gene module associated with the antibiotic resistance phenotypes in *P. aeruginosa* **(Langfelder & Horvath, 2008).** The analysis pipeline started with data preprocessing, where the gene presence-absence matrix and strain metadata were imported, and strains were kept for further analysis if they were present in both data sets. To guarantee the presence of adequate variation for network building, genes were filtered based on their prevalence, retaining those present in 5% to 95% of the strains **(Ikhimiukor et al., 2025).** This eliminated genes that were universally present, as well as those that were very rare and would only add noise to the data rather than providing relevant co-occurrence information. The final data set consisted of strains with known phenotypes for resistance and susceptibility, which were represented as a binary trait for later correlation studies.

#### 2.10.2 Network Construction

Network building followed standard WGCNA methods **(Langfelder & Horvath, 2008).** We used a suitable soft threshold power to build a scale-free topology and checked the scale-free topology fitting index and the mean connectivity values for the soft threshold power values ranging from 1 to 20. A soft threshold power of 6 was used, as recommended when linear relationships are not assumed **(Zhang & Horvath, 2005).** Using this capacity, we were able to compute unsigned adjacencies for all gene pairs and then transform these data into topological overlap matrices (TOM). These matrices allow for the determination of direct and indirect connections **(Langfelder & Horvath, 2008).** We also performed hierarchical clustering on the dissimilarity matrix based on the topological overlap matrix data using average linkage. To define the module membership, we initially applied the method of dynamic tree cutting with a module size of at least 5 genes **(Yu et al., 2011).** To further refine the module membership, we computed the module eigengenes, the first principal component of the expression for each module, and then merged the modules based on eigengene correlation values of 0.75 or higher, a dissimilarity value of 0.25 (https://support.bioconductor.org/p/124692/).

#### 2.10.3 Module-Trait Association & Hub Gene Identification

Module trait relationships were established by correlating the module eigengenes with the resistance phenotype. Statistical significance was established using Student’s correlation p values, with p < 0.05 being the threshold for significance **(Zhang et al., 2019).** Hub genes were identified in the significant modules by calculating the module membership (correlation of gene presence-absence profiles with the module eigengene) and gene significance (correlation of gene presence-absence profiles with the phenotype) **(Zhang et al., 2022).** Hub score is the product of absolute values of module membership and gene significance. Hub genes were selected based on the top ten genes in each significant module, provided that they had a non-zero hub score. However, modules with fewer than ten genes or fewer than three genes were excluded.

All WGCNA analyses were performed using R (v4.5.2) within the RStudio environment (https://www.r-project.org/).

### 2.11 Comparative Network Topology Analysis

#### 2.11.1 Network Reconstruction

In order to systematically elucidate the architectural differences between the resistance associated and susceptible gene co-occurrence networks, a topological analysis of the networks was carried out using the WGCNA approach for module detection (**Langfelder & Horvath, 2008**). For this purpose, the reconstruction of the networks was performed separately using the resistant and susceptible strain populations using the same presence/absence matrix as used in the WGCNA approach. The edge weights were calculated using the Spearman rank correlation between the two genes, and the threshold used for the correlation was ρ ≥ 0.3, together with the correction for multiple testing using the FDR < 0.05 threshold (**Johansson et al., 2011; Randhawa & Acharya, 2015**). This resulted in the construction of two networks: one representing the resistant and the other the susceptible strain populations.

#### 2.11.2 Network Degree Distribution Analysis

To further understand the global architecture of the gene co-occurrence networks across the globe, the degree distribution of the resistant strain networks was compared to the degree distribution of the susceptible strain networks **(Barabási & Oltvai, 2004).** For each gene node, the degree was represented as the number of connections formed with other genes, and the data was collected from the results of the network centrality calculation performed. Frequency histograms were prepared for the resistant and susceptible strain networks, each divided into 50 equal intervals over the range from 0 to the maximum degree value observed in the networks for the two phenotypes **(Burke et al., 1988).** Due to the right-skewed nature of the degree distribution, the y-axis scale was set to a logarithmic scale for better visualization of the data. In addition to the graphical representation of the data, the average degree for each phenotype was also calculated as a simple measure for the quantitative comparison of the global connectivity of the resistant and susceptible strain networks **(Jia et al., 2025).**

#### 2.11.3 Global Topology Metrics

Global network topology was also characterized by three standard parameters calculated for each of these networks: density of the network (fraction of edges that can exist actually exist), average clustering coefficient (tendency of nodes to cluster together), and transitivity (closed triplets vs. open triplets ratio) **(Bedru et al., 2020; Dekker et al., 2019; Nesterov, 2024).** These parameters together will allow us to determine if resistant strains have a different network organization compared to susceptible strains, namely if it is more or less cohesive, clustered, or hierarchical.

#### 2.11.4 Module Connectivity Comparison

To compare module-level connectivity between phenotypes, intramodular connectivity values, kWithin, were obtained from the WGCNA output and stratified by resistant and susceptible strains **(Sun et al., 2025).** For each module, the average connectivity was computed within each set of strains, and the comparison was performed using the Wilcoxon rank sum test **(Fang et al., 2012).** For the primary comparison and better visualization, we chose to examine these six modules that had the highest average connectivity in the resistant network. Finally, we compared the average kWithin values of these modules between phenotypes to see if modules that had higher connectivity in resistant strains also had higher connectivity in susceptible strains or vice versa **(Peng et al., 2021).**

In addition, a specific analysis was carried out on three modules: darkred, plum3, and grey, which included genes that had overlap between the WGCNA and GWAS studies. The same approach as described above was used in this analysis, although the selection criteria were not based on the highest connectivity in the resistant strains. Instead, these modules were specifically selected due to their biological importance. This analysis provided us the opportunity to evaluate whether the modules that contained genes supported by the GWAS studies maintained their connectivity patterns or had differences that were strain-specific in relation to antibiotic resistance. The overall methodology is illustrated in Figure Schematic 1.

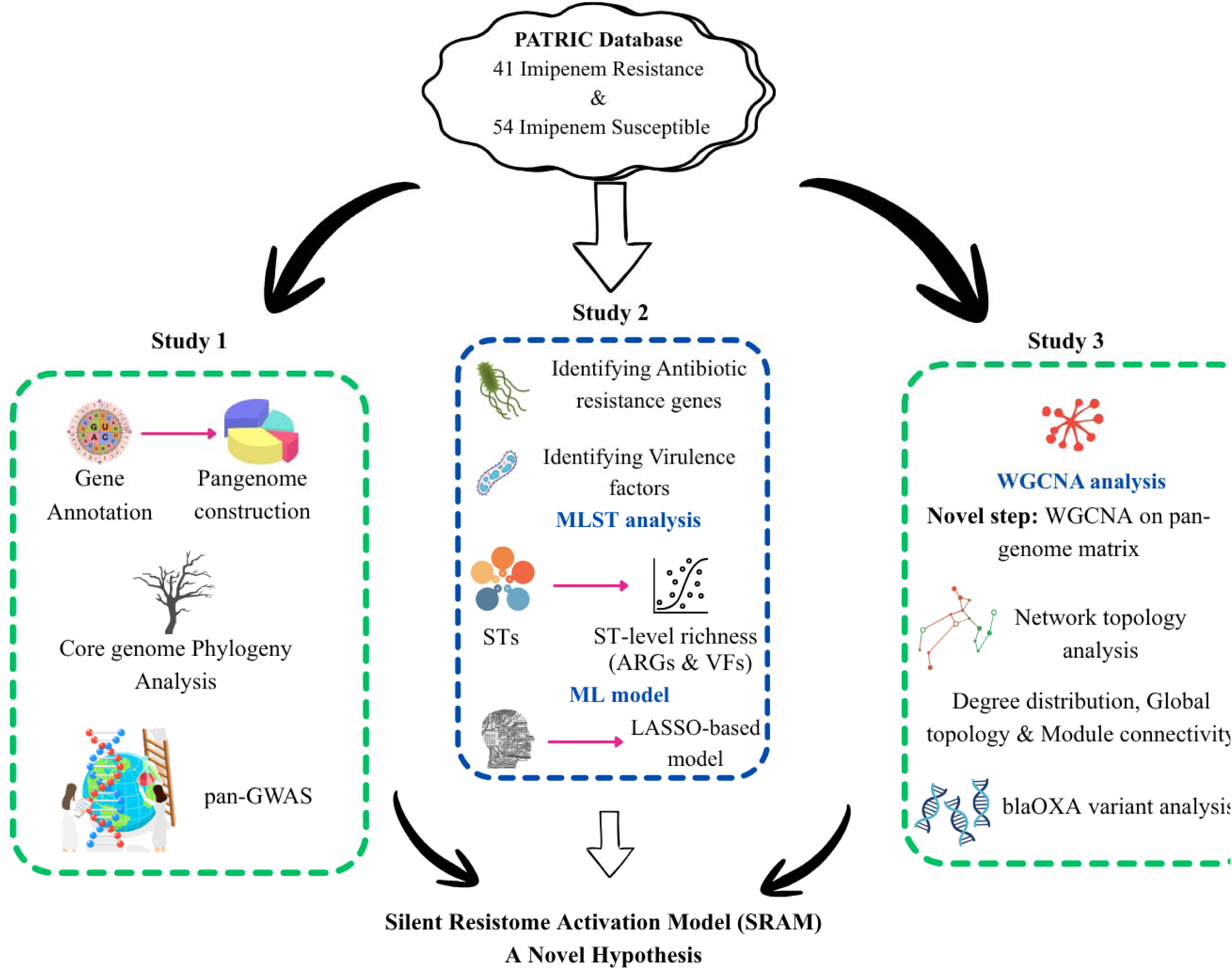

**Schematic 1**: Overview of the analytical workflow used to study 95 different *P. aeruginosa* strains, including both imipenem-resistant and imipenem-susceptible isolates. The schematic illustrates the major steps of the study from sample selection to genomic analyses. *The figure was created using an online designing tool, Canva* (https://www.canva.com/).

## 3. Results and Discussion

### 3.1 Characteristics of the *P. aeruginosa* Isolate Collection

The study examined a total of 95 *P. aeruginosa* isolates, with 41 being resistant to Imipenem and 54 being susceptible to Imipenem. The majority of the *P. aeruginosa* isolates were obtained from the U.S. Food and Drug Administration (FDA) and the Centers for Disease Control and Prevention (CDC), with some being obtained from the Multidrug-Resistant Organism Repository and Surveillance Network (MRSN). In this study, we have only considered those samples for which MIC > 64 mg/L for resistant strains and MIC < 2 mg/L for susceptible strains. We have intentionally avoided those strains for which MIC values were intermediate, i.e., 3–63 mg/L, as these would have created phenotypic ambiguity in our results. This strict selection criterion was followed so that we could identify robust genomic differences associated with imipenem resistance. Quality metrics for genome assembly, including completeness, contamination, contig number, N50, L50, GC content, etc., are provided in Supplementary File 1.

### 3.2 Identification of Antibiotic Resistance Genes in Imipenem-Resistant Isolates

Antibiotic resistance genes were identified in all 41 imipenem-resistant *P. aeruginosa* isolates using AMRFinderPlus. Although some isolates carried multiple allelic variants within the same gene family, results were summarised at the gene-family level to provide a clear overview. As shown in the **Figure 1A**, the *bla*OXA and *bla*PDC genes were detected in all resistant isolates. In addition, the resistance genes *aph*, *catB*, and *fosA* were present in nearly 90% of the strains. Several other resistance determinants were also observed, including *bla*VIM, which was identified in 13 of the 41 isolates (**Figure 2A**). This gene encodes a metallo-β-lactamase that is well known for its role in carbapenem resistance. While *bla*PDC is not typically associated with carbapenem hydrolysis, certain variants of *bla*OXA are recognised for their carbapenemase activity.

### 3.3 Identification of resistance determinants in imipenem-susceptible isolates

The same AMRFinderPlus analysis was applied to the 54 imipenem-susceptible *P. aeruginosa* isolates. As shown in **Figure 1B**, both *bla*PDC and *bla*OXA genes were detected in all susceptible genomes. While *bla*PDC is not known to confer carbapenem resistance, it was unexpected to observe that certain *bla*OXA variants, which have been previously associated with carbapenemase activity, were also present in the susceptible group. It seems that for the activation of antibiotic resistance genes, some accessory genes are required. For example, when we backtracked the results obtained from AMRFinderPlus, several blaOXA variants already known to hydrolyze carbapenem antibiotics, such as blaOXA-2, blaOXA-486, and others that have been well characterized in earlier studies, were detected in both imipenem-resistant and imipenem-susceptible isolates **(Antunes et al., 2014; Dale et al., 1985; Zaidi et al., 2023).** While their presence in resistant strains is expected, their detection in susceptible isolates was unexpected and noteworthy. Results of all the 95 samples of AMRFinderPlus are provided in Supplementary File 2.

In addition, the resistance genes *aph*, *catB*, and *fosA* were identified in 53 of the 54 isolates (98.1%), indicating that these markers are nearly ubiquitous among imipenem-susceptible strains. Together, these observations highlight the importance of directly comparing resistant and susceptible isolates to better understand the genetic features that truly distinguish their resistance profiles.

**Figure 1:**
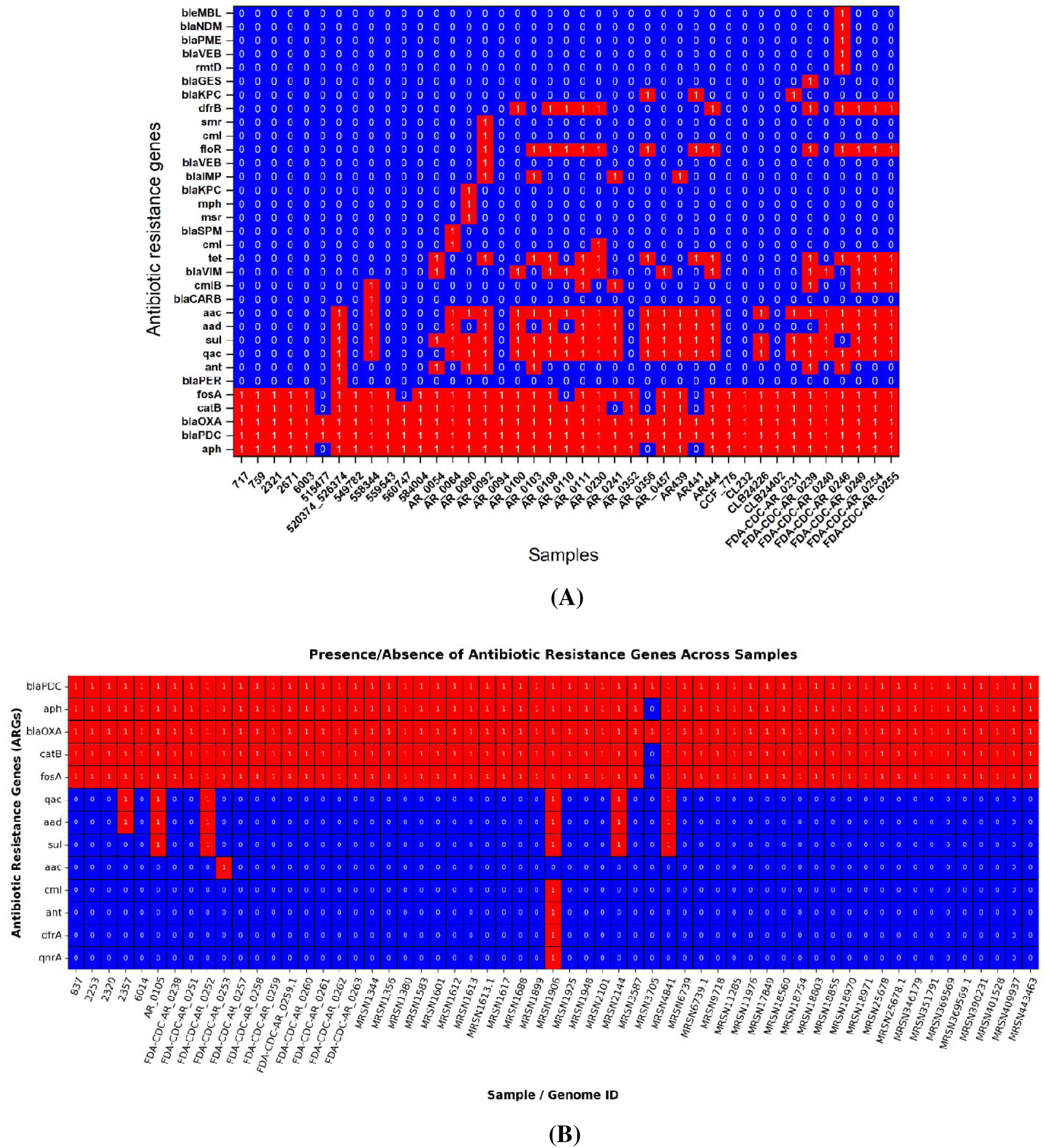
Binary presence–absence matrix illustrating antibiotic resistance genes in *P. aeruginosa* isolates. In the heatmap, blue (0) indicates the absence of a gene, while red (1) indicates its presence. Figure **(A)** represents imipenem-resistant strains, and figure **(B)** represents imipenem-susceptible strains. *Figure (A) was generated using Origin 2025b* (https://www.originlab.com/)*, whereas Figure (B) was produced using Python-based libraries*.

### 3.4 Comparison of Antibiotic Resistance Genes Between Imipenem-Resistant and Imipenem-Susceptible Isolates

As presented in **Figure 2**, the presence-absence analysis of the antibiotic resistance genes showed that the imipenem-resistant group had significantly more diversity of resistance determinants compared to the susceptible group. For example, the resistant group showed diverse β-lactamase gene families. A total of 15 different β-lactamase family genes were found in the resistant group, compared to only two different β-lactamase family genes found in the imipenem-susceptible group.

Some other metallo-beta-lactamase genes, such as blaVIM, blaIMP, and blaNDM, were found specifically within the resistant group. These genes are often linked to mobile genetic elements such as plasmids **(Boyd et al., 2020).** When the total number of resistance gene diversity was examined, it was found that the imipenem-resistant group possessed 33 different antibiotic resistance gene families, whereas the susceptible group possessed only 13. These findings indicate that imipenem resistance is not likely influenced by a single gene, but rather the culmination of multiple genes.

**Figure 2:**
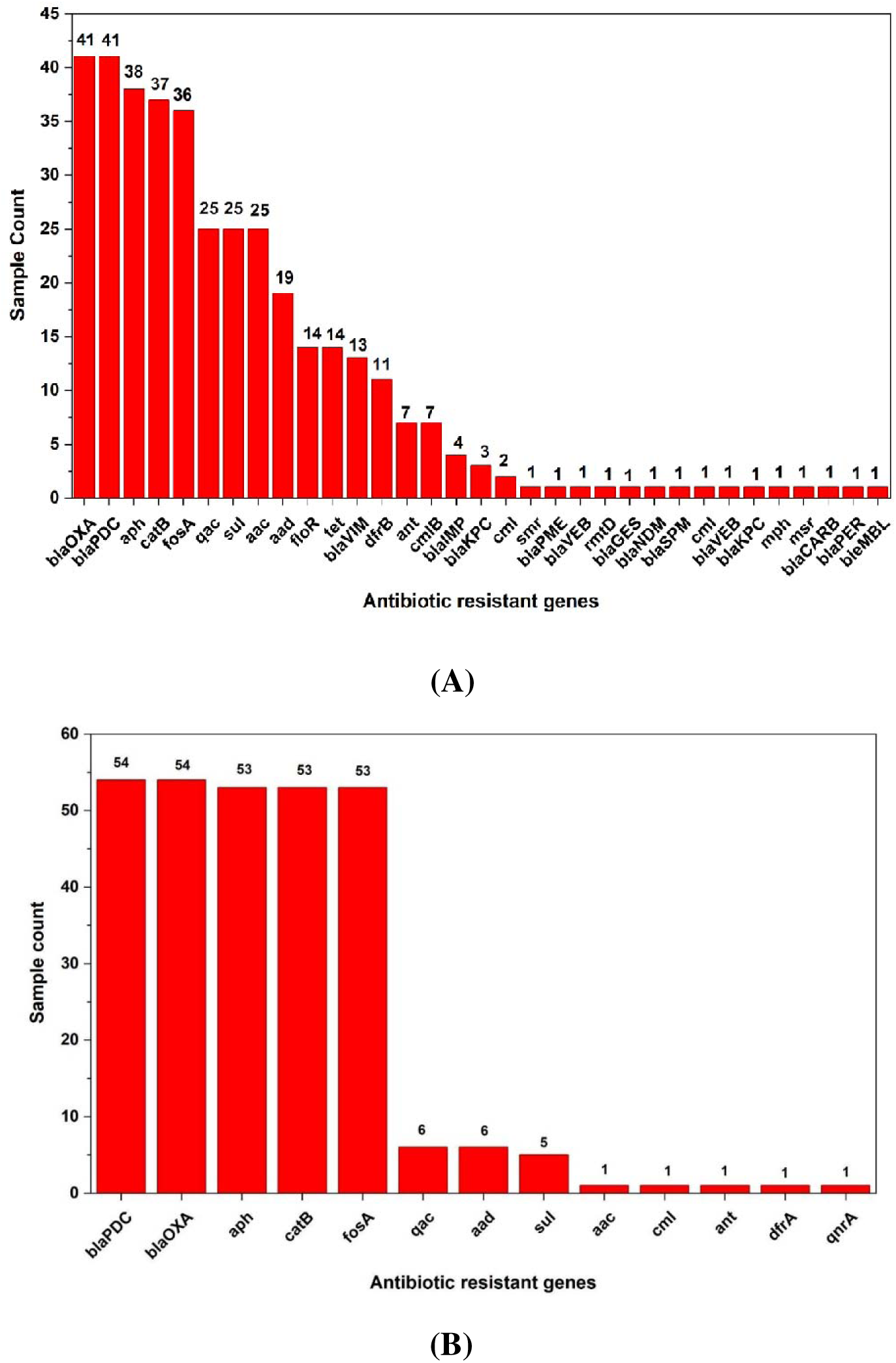
Column bar plots showing the richness of antibiotic resistance genes in *P. aeruginosa* isolates. Figure **(A)** represents imipenem-resistant strains, while Figure **(B)** represents imipenem-susceptible strains. *Both the figures were generated using Origin 2025b* (https://www.originlab.com/).

### 3.5 Variant Analysis of blaOXA

Variant-level analysis of the blaOXA gene family was carried out by backtracking the results obtained from AMRFinderPlus. Across all isolates, we identified a total of 28 distinct blaOXA variants. To better understand their functional relevance, we conducted an extensive literature review to determine whether each variant had previously been reported, whether it showed carbapenemase activity or not. As summarized in **Table 1**, no published information was available for 12 of the 28 identified variants, indicating that these variants have not yet been described in earlier studies. This highlights a substantial gap in current knowledge regarding the blaOXA variant landscape.

A particularly noteworthy example is shown in **Figure 1A**. One imipenem-resistant isolate, strain 515477, carried only two antibiotic resistance gene families: blaOXA and blaPDC. Since blaPDC is not known to confer carbapenem resistance, this suggests that resistance in this strain is most likely driven by the blaOXA gene. Further back-tracing revealed that the variant present in this isolate was blaOXA-1284. To date, this variant has not been reported in the literature, and its ability to hydrolyze carbapenems remains unknown. It emphasizes the need for focused functional studies on uncharacterized blaOXA variants to better understand their potential role in carbapenem resistance.

Another notable observation was that several blaOXA variants already known to hydrolyze carbapenem antibiotics, such as blaOXA-2, blaOXA-486, and others that have been well characterized in earlier studies, were detected in both imipenem-resistant and imipenem-susceptible isolates. While their presence in resistant strains is expected, their detection in susceptible isolates was unexpected and noteworthy.

The presence of these carbapenemase-associated variants in susceptible strains suggests that gene presence alone is not sufficient to confer resistance. Instead, resistance likely depends on additional factors, including the expression level of these genes, their genomic context, and the involvement of other regulatory or accessory pathways. In susceptible isolates, these blaOXA variants may remain functionally silent under current conditions, yet their genomic positioning could allow rapid activation or enhanced expression when exposed to antibiotic pressure.

This finding highlights the existence of a “silent resistome,” where resistance genes are present but not phenotypically expressed. It also emphasizes the importance of strategies aimed at preventing resistance emergence rather than responding only after resistance has developed.

**Table 1:** Distribution of different *bla*OXA variants among *P. aeruginosa* isolates. The table shows variants found exclusively in imipenem-resistant strains **(R)**, exclusively in imipenem-susceptible strains **(S)**, or present in both groups **(B).**

| OXA-Variant | Carbapenemase activity | Reference |
| --- | --- | --- |
| blaOXA-10 (B) | Weak activity | Maurya et al., 2017; Alonso-García et al., 2023 |
| blaOXA-101 (R) | No known activity | Porto et al., 2010; Kang & Cui, 2025 |
| blaOXA-1028 (B) | Known activity | AbdulHak et al., 2025 |
| blaOXA-1127 (S) | No information | No information |
| blaOXA-1131 (S) | No information | No information |
| blaOXA-1135 (S) | No information | No information |
| blaOXA-1174 (S) | No information | No information |
| blaOXA-1254 (S) | No information | No information |
| blaOXA-1267 (S) | No information | No information |
| blaOXA-1275 (S) | No information | No information |
| blaOXA-1284 (R) | No information | No information |
| blaOXA-1290 (R) | No information | No information |
| blaOXA-1293 (S) | No information | No information |
| blaOXA-2 (B) | Known activity | Antunes et al., 2014; Dale et al., 1985 |
| blaOXA-395 (B) | Known activity | Bai et al., 2024 |
| blaOXA-396 (B) | Weak activity | Mack et al., 2025; Tsilipounidaki et al., 2024 |
| blaOXA-4 (R) | Known activity | Kingsley & Verghese, 2013 |
| blaOXA-486 (B) | Known activity | Zaidi et al., 2023 |
| blaOXA-488 (B) | Known activity | Streling et al., 2022 |
| blaOXA-494 (B) | Known activity | Zaidi et al., 2023 |
| blaOXA-50 (B) | Weak activity | Girlich et al., 2004 |
| blaOXA-56 (R) | Weak activity | Porto et al., 2010; Galetti et al., 2019 |
| blaOXA-846 (R) | Known activity | Mondol et al., 2024 |
| blaOXA-847 (B) | No known activity | Kang & Cui, 2025 |
| blaOXA-9 (R) | No known activity | Meng et al., 2023 |
| blaOXA-904 (R) | Known activity | AbdulHak et al., 2025 |
| blaOXA-914 (S) | No information | No information |
| blaOXA-937 (S) | No information | No information |

### 3.6 Pan-genome Analysis and Core Genome Phylogeny Construction

We carried out a pan-genome analysis for the 41 *P. aeruginosa* isolates that were resistant to Imipenem. As demonstrated in **Figure 3A**, we were able to identify 12,988 gene families that were unique. Of these gene families, only 34% were part of the core genome, while 44.7% were cloud genes. This indicates that there was diversity among the resistant isolates.

We also performed the same analysis on the 54 imipenem susceptible isolates, and the results are shown in **Figure 3B**. What is interesting is that this group had 11,984 gene families, which is almost 1,000 fewer than the resistant group, although this group had more isolates than the resistant group. Normally, a larger number of isolates should correspond to a larger number of gene families. However, the reverse trend in this case suggests that the imipenem-resistant isolates have a more diverse gene repertoire than the susceptible isolates.

In order to have a broader overview of the results, we also constructed the pan-genome of the 95 isolates taken together (resistant + susceptible) as shown in **Figure 3C**. We found that 56.3% of the gene families fell within the cloud genome fraction, again highlighting the high genomic variability of *P. aeruginosa*.

A phylogenetic tree was constructed based on the core genome, and we analyzed the evolutionary relationships between these strains. As shown in **Figure 3D**, no obvious clusters are observed that distinguish imipenem-resistant strains from imipenem-susceptible strains. These results indicate that imipenem resistance does not seem to emerge from a dominant resistant clone. Rather, imipenem resistance appears to emerge independently from multiple strains, probably due to acquisition of additional genetic elements. A tree map of rectangles was provided as shown in Supplementary File 3.

This is consistent with the known characteristics of *P. aeruginosa*, as carbapenem resistance in this species is usually the result of a combination of events involving horizontal gene transfer and independent chromosomal mutations in the efflux mechanisms, porins, and beta-lactamases.

This leads to a mosaic distribution of resistance traits that does not strictly follow the core genome phylogeny. These data suggest that imipenem resistance in *P. aeruginosa* is a complex and multifactorial phenomenon that has evolved independently multiple times from a variety of genetic backgrounds, rather than resulting from the spread of a single resistant lineage.

**Figure 3:**
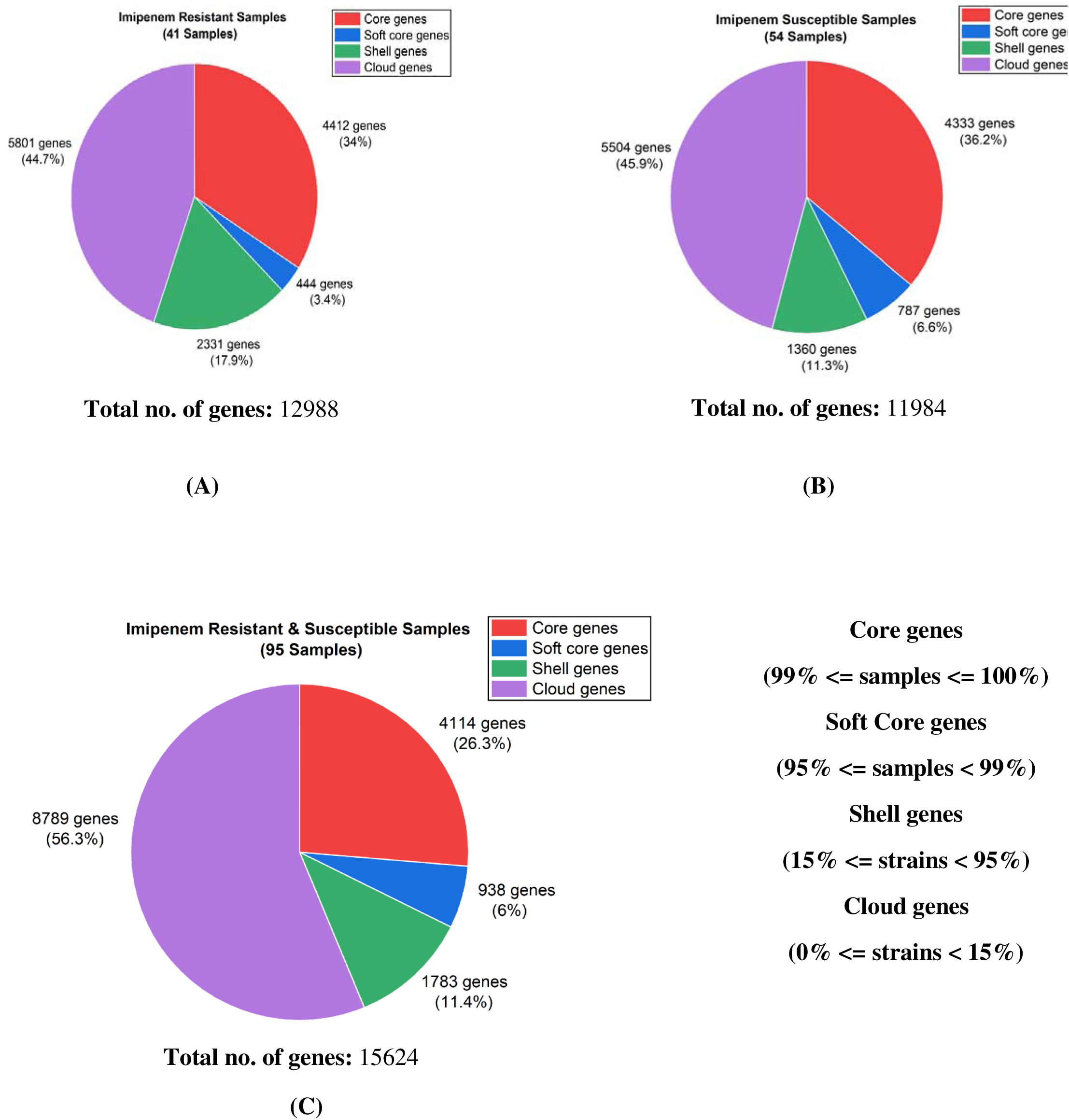
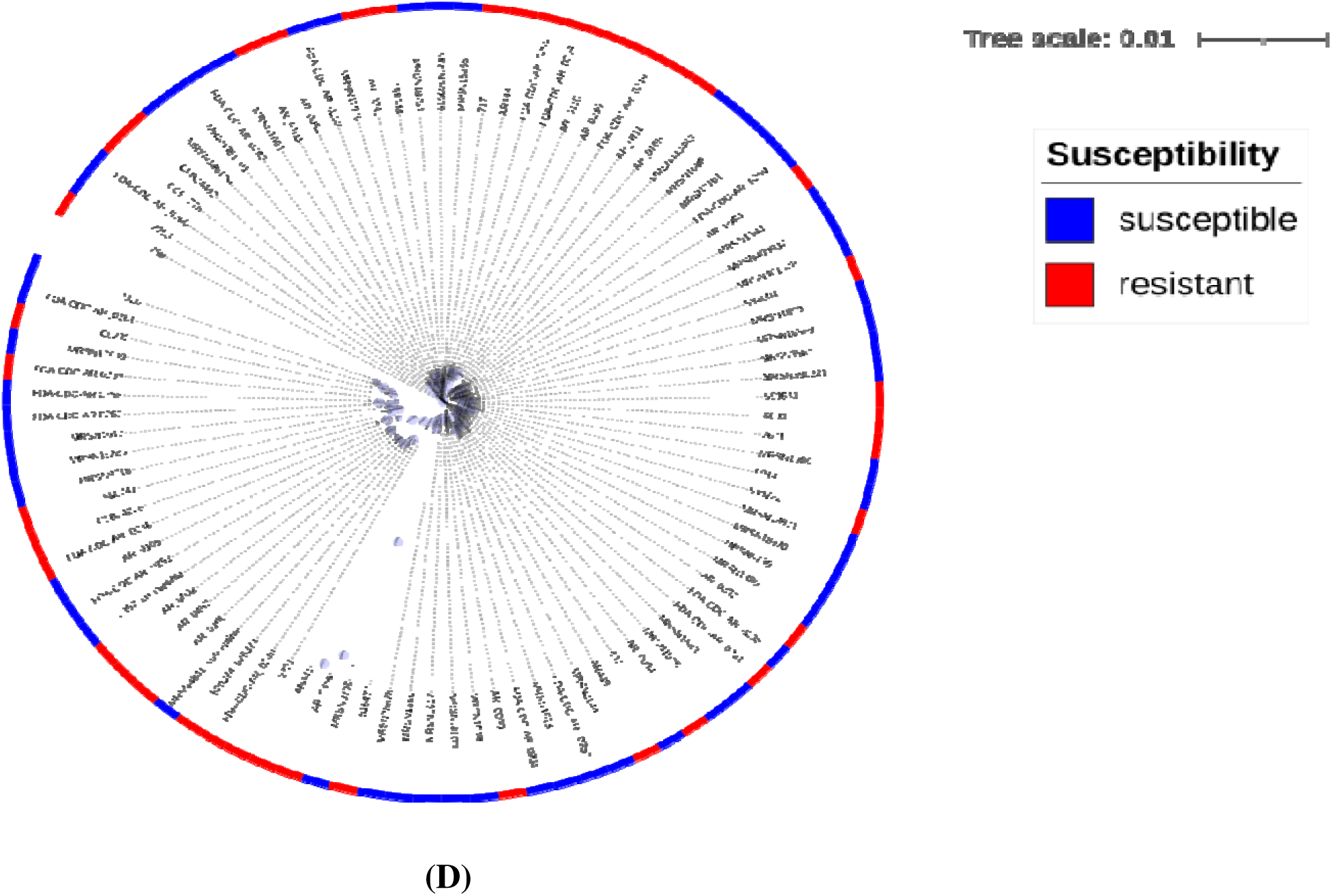
Pan-genome and core-genome phylogenetic analysis of *P. aeruginosa* strains. Figure **(A)** shows the pan-genome of imipenem-resistant isolates, figure **(B)** shows the pan-genome of imipenem-susceptible isolates, and figure **(C)** shows the combined pan-genome of both groups. Figure presents the core-genome phylogenetic tree. *Figures **(A–C)** were generated using Origin 2025b (*https://www.originlab.com/*), while figure **(D)** was created using the iTOL web server (*https://itol.embl.de/).

### 3.7 Pan-GWAS Analysis

To carry out the pan-genome-wide association study, we used two different approaches, Scoary and PySEER, and found genes that are significantly associated with the imipenem-resistant phenotype.

#### 3.7.1 Scoary Result

Using Scoary, we found that there are 250 genes that are significantly associated with imipenem resistance, as shown in **Figure 4A**, considering the criteria of the adjusted p-value by Benjamini and Hochberg to be less than 0.05 and the odds ratio to be more than 1, showing that the genes are positively associated with the resistant phenotype.

Out of these 250 genes, 99 had an infinite odds ratio. This means that these genes are present in all imipenem-resistant strains or are absent from imipenem-susceptible strains. Thus, these genes are perfectly correlated with imipenem resistance. Since the x-axis of the volcano plot in **Figure 4B** shows the odds ratios, genes with infinite odds ratios could not be included in this plot. However, all other genes are included in this plot, giving a general overview of their associations.

To further evaluate the usefulness of these genes as possible resistance markers, their sensitivity and specificity were calculated. The results are depicted in **Figure 4C**. In some instances, different genes had similar sensitivity and specificity values, which resulted in similar points being represented in the figure. Genes that were more specific were less likely to be found in imipenem-susceptible isolates, while those that were more sensitive were more likely to be found in resistant isolates.

These results, therefore, provide a way of assessing how well each of these genes can differentiate between resistant and susceptible strains. A complete list of the 250 genes identified by Scoary, along with their statistical details, is shown in Supplementary File 4.

**Figure 4:**
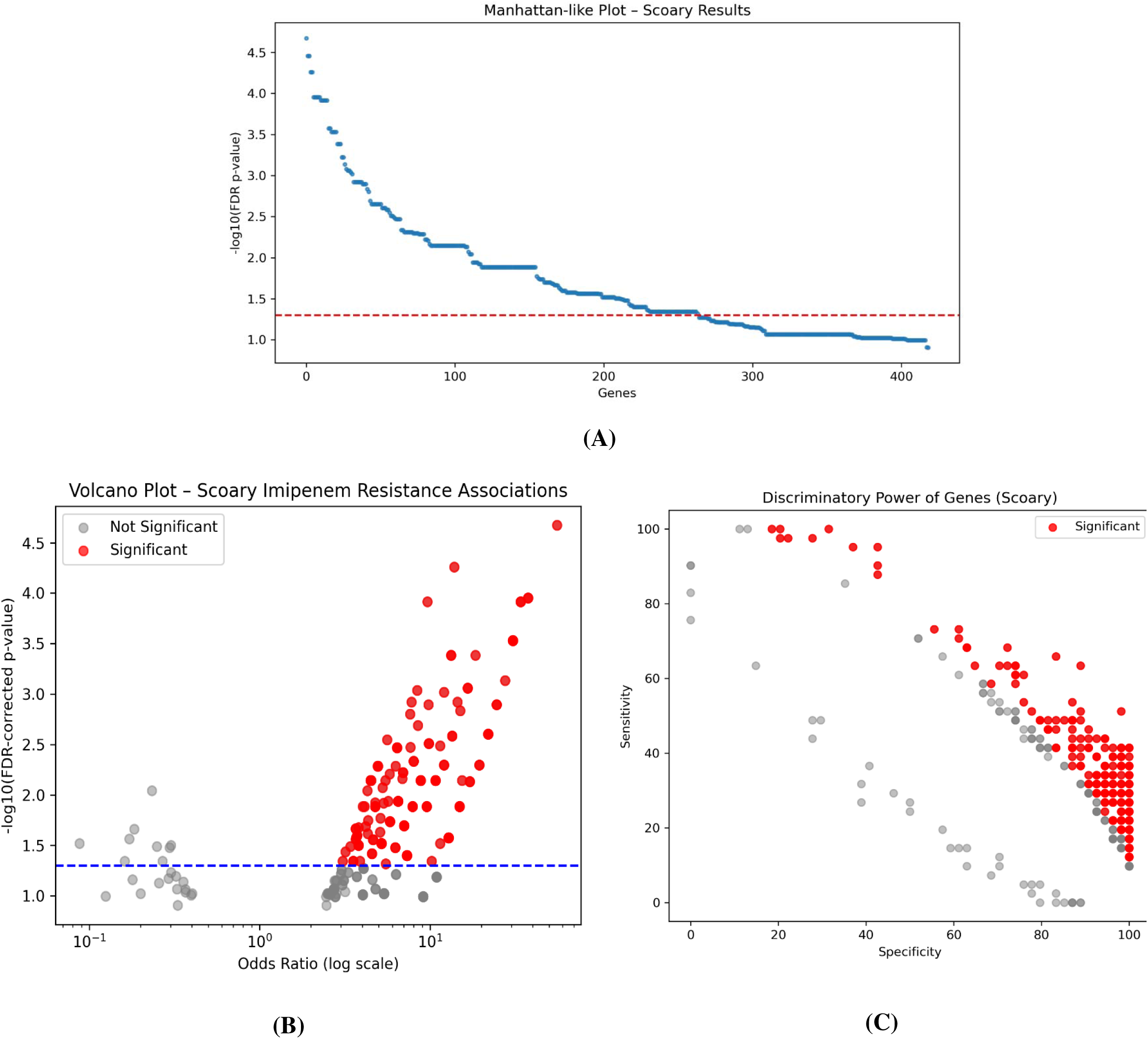
Output of the Scoary pan-GWAS analysis. **Figure (A)** shows a Manhattan-like plot, where the red horizontal line marks the Benjamini–Hochberg adjusted p-value cutoff of 0.05; 250 genes exceed this threshold. **Figure (B)** presents a volcano plot, with significant genes in red bubbles. **Figure (C)** shows a scatter plot illustrating the sensitivity and specificity of significant genes, with red points indicating genes with p-values below 0.05. *All plots were generated using Python-based libraries*.

#### 3.7.2 PySEER Result

To enhance the robustness of our results, we again carried out the pan-GWAS analysis using the PySEER approach. This analysis revealed that there are 26 gene families that are statistically significantly associated with the imipenem-resistant phenotype, i.e., LRT adjusted p-values < 0.05 after correction for multiple testing. As shown in **Figure 5**, it was found that 25 genes had positive beta coefficients, implying that the presence of these genes was positively correlated with the imipenem-resistant phenotype. Conversely, it was found that one gene had a negative beta coefficient, implying that the presence of this gene was negatively correlated with the imipenem-resistant phenotype.

The list of all the genes identified using the PySEER approach, along with all the statistical parameters, is provided in Supplementary File 5.

**Figure 5:**
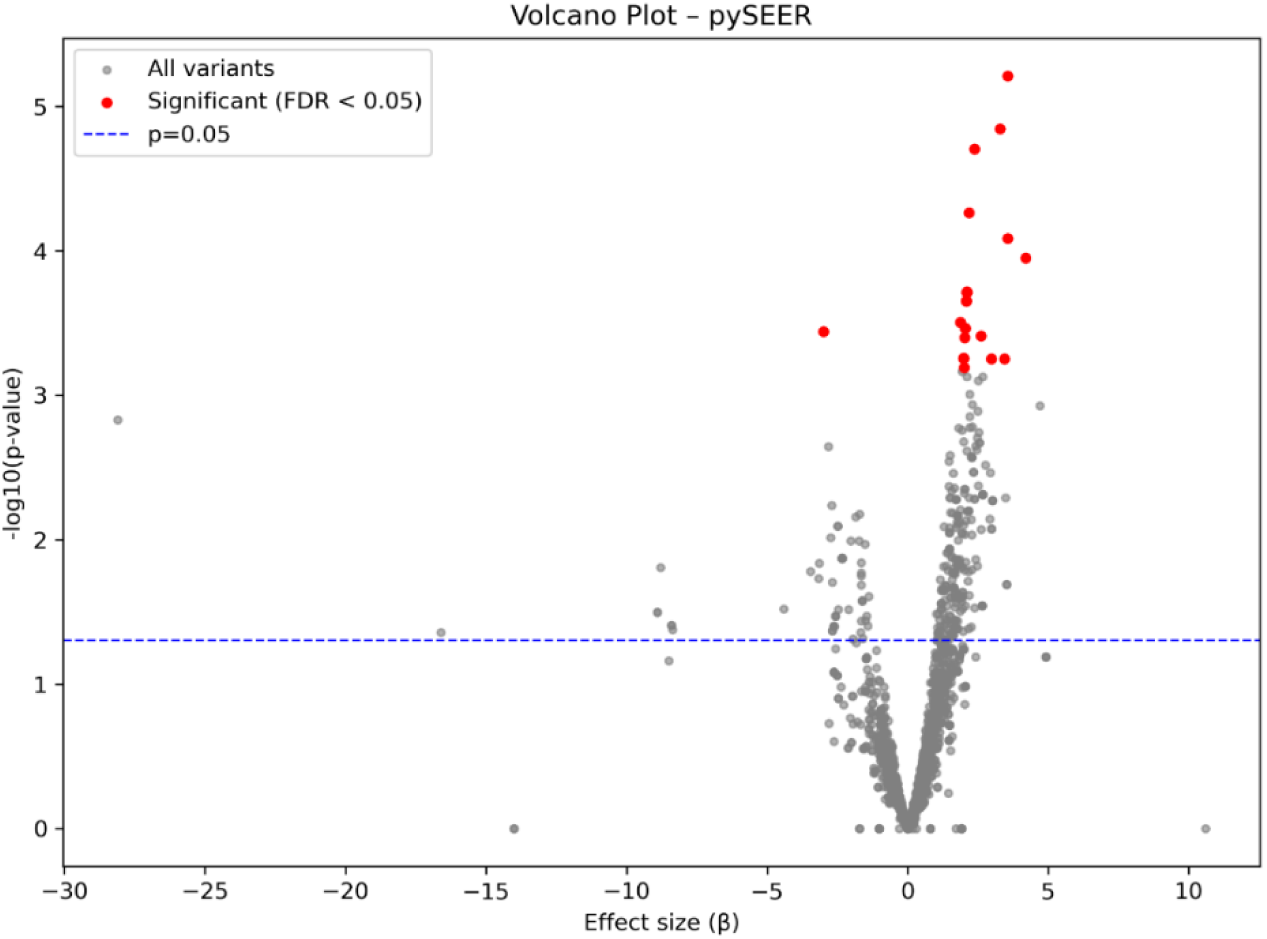
Results of the PySEER pan-GWAS analysis. The volcano plot highlights genes significantly associated with imipenem resistance, shown as red points. *The figure was generated using Python-based libraries*.

#### 3.7.3 Overlapping the Scoary and PySEER results

To determine the best and most robust associated genes with resistance, the results obtained from the analyses by Scoary and Pyseer were compared. From the results shown in **Figure 6A** and **Figure 6B**, Scoary identified 250 significant genes, while Pyseer identified 26 genes associated with imipenem resistance. When the results obtained from the two analyses were overlapped, it was determined that the genes had 12 in common. The genes have been determined to be the best and most associated with the subject, as they were obtained from two analyses.

The list of these 12 genes, along with their corresponding statistical parameters, is provided in **Table 2**.

**Figure 6:**
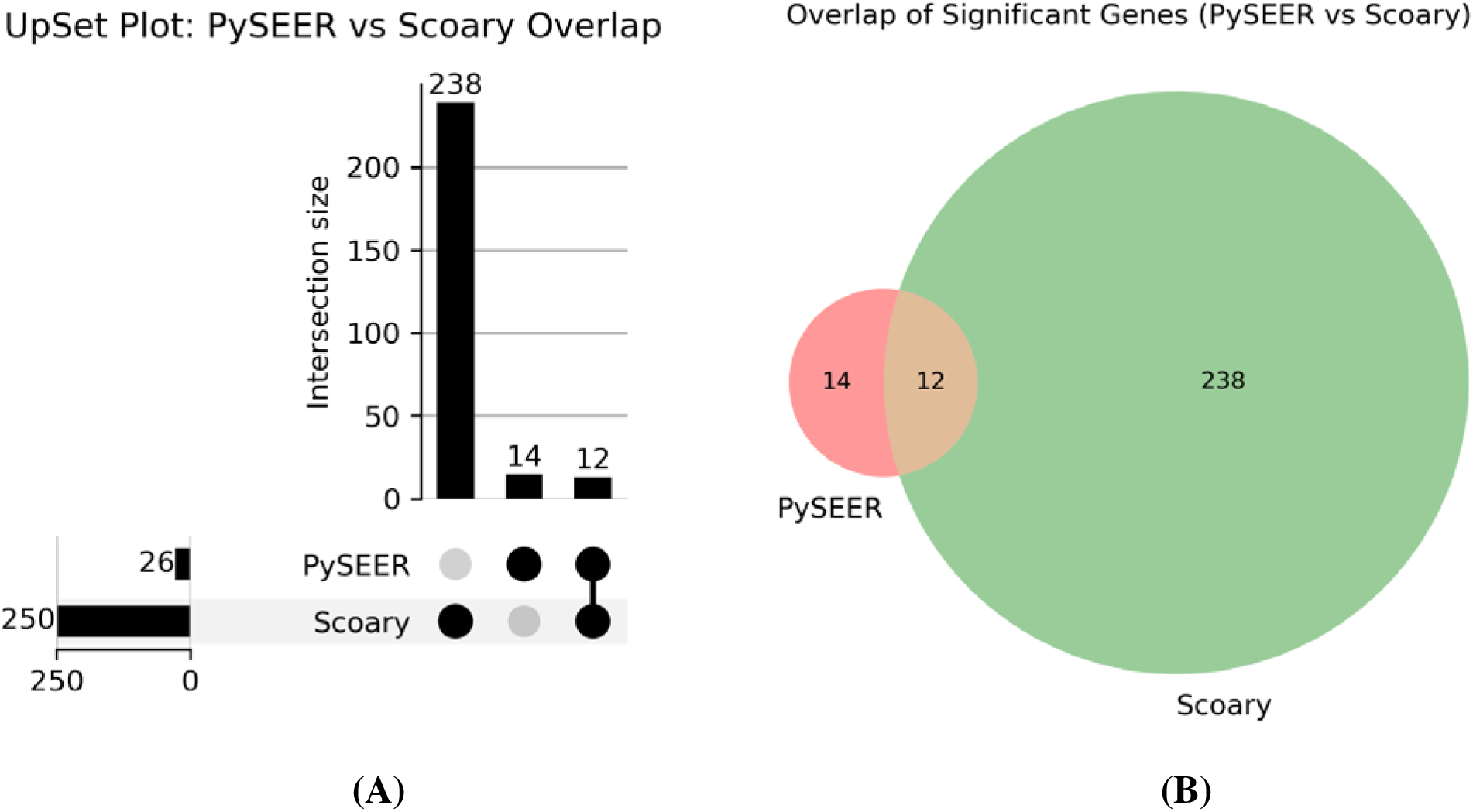
Overlap of genes identified by Scoary and PySEER. Figure **(A)** shows an UpSet plot illustrating genes shared between both tools as well as genes uniquely detected by each tool. Figure **(B)** presents a Venn diagram summarizing the number of common and tool-specific genes identified in the analysis. *Both the figures were generated using Python-based libraries*.

**Table 2:**
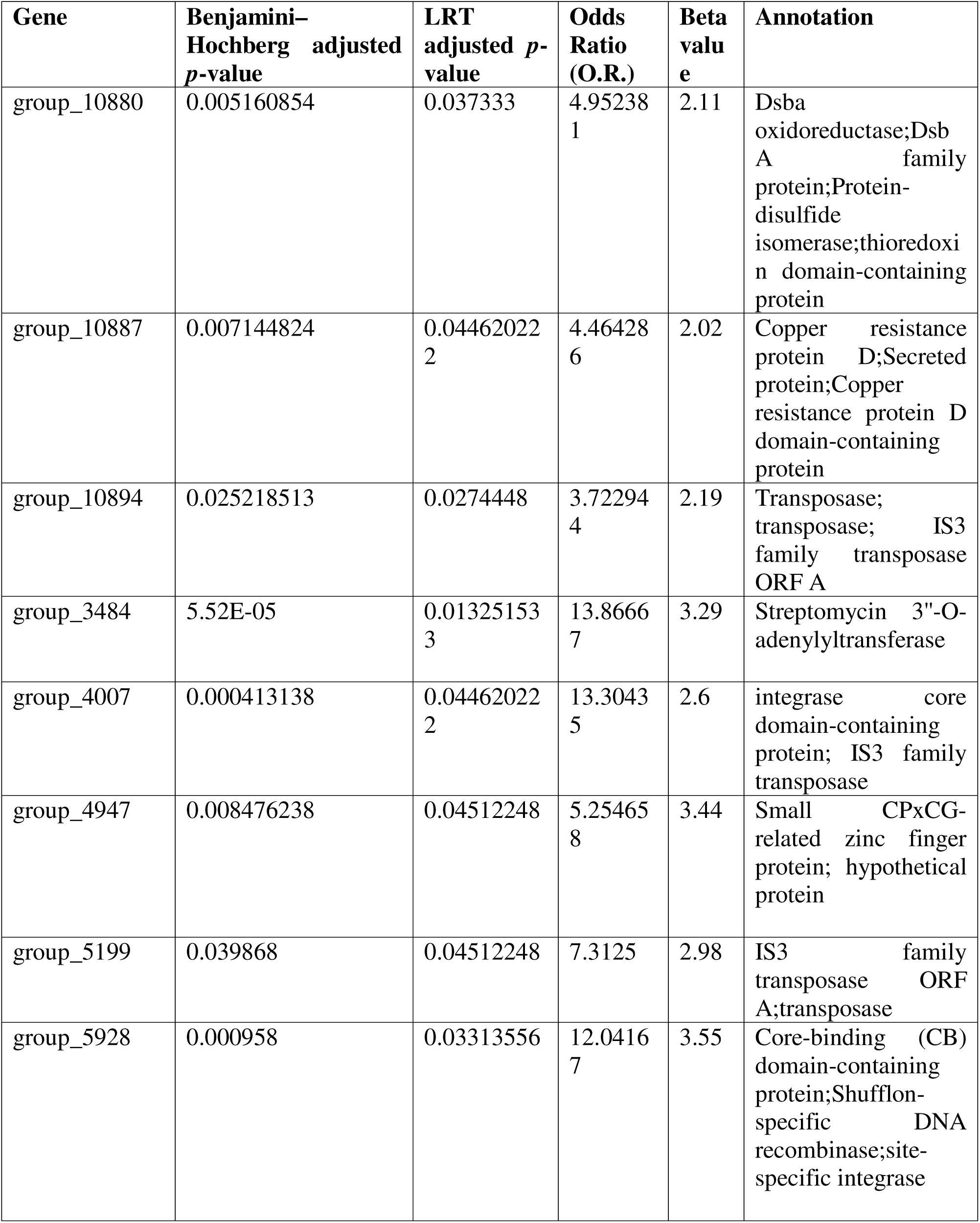
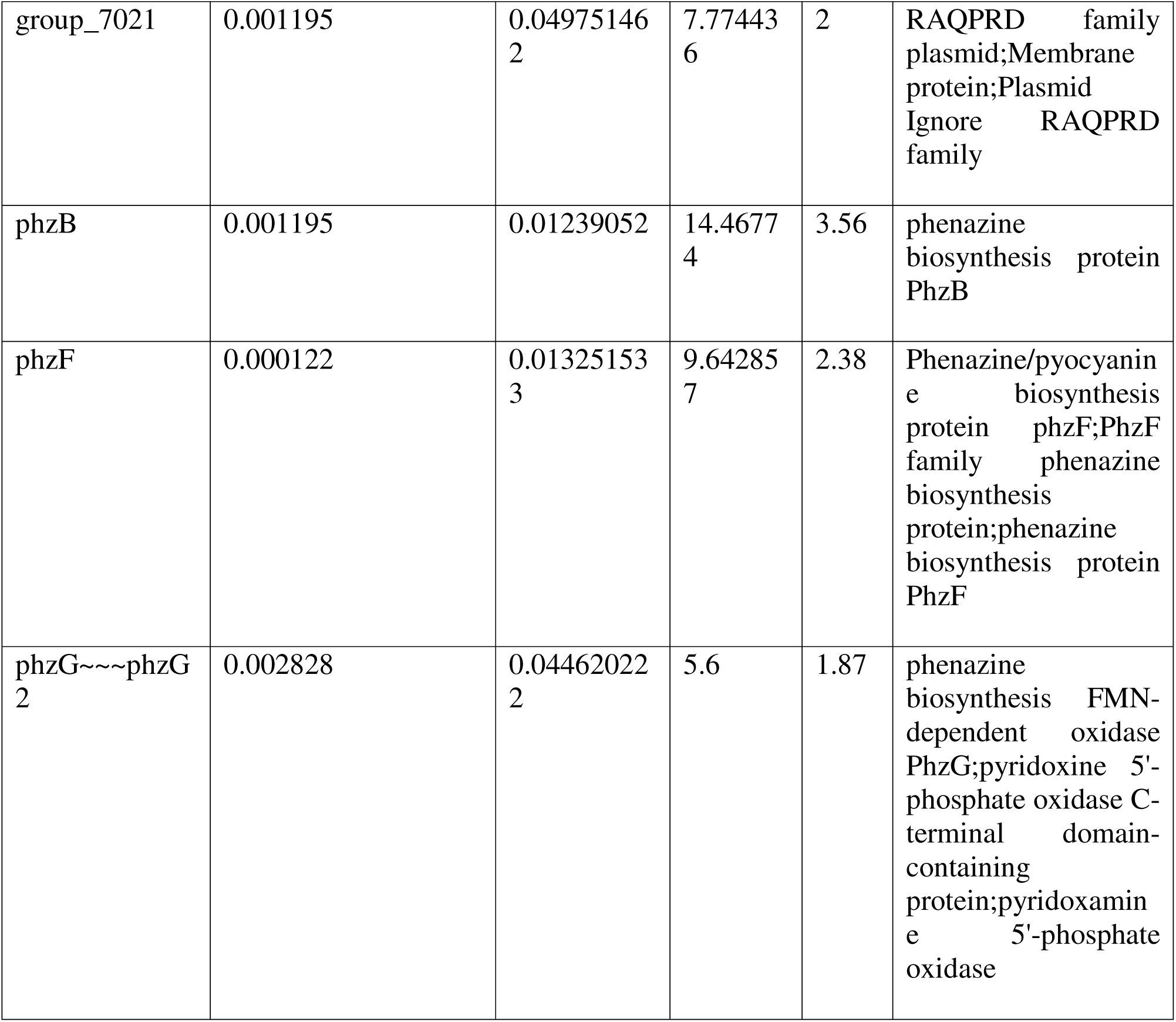
List of genes commonly identified by both Scoary and PySEER, along with their corresponding statistical parameters.

| Gene | Benjamini–Hochberg adjusted *p*-value | LRT adjusted *p*-value | Odds Ratio (O.R.) | Beta value | Annotation |
| --- | --- | --- | --- | --- | --- |
| group_10880 | 0.005160854 | 0.037333 | 4.952381 | 2.11 | Dsba oxidoreductase;Dsb A family protein;Protein-disulfide isomerase;thioredoxin domain-containing protein |
| group_10887 | 0.007144824 | 0.04462022 | 4.464286 | 2.02 | Copper resistance protein D;Secreted protein;Copper resistance protein D domain-containing protein |
| group_10894 | 0.025218513 | 0.0274448 | 3.722944 | 2.19 | Transposase; transposase; IS3 family transposase ORF A |
| group_3484 | 5.52E-05 | 0.01325153 | 13.86667 | 3.29 | Streptomycin 3"-O-adenylyltransferase |
| group_4007 | 0.000413138 | 0.04462022 | 13.30435 | 2.6 | integrase core domain-containing protein; IS3 family transposase |
| group_4947 | 0.008476238 | 0.04512248 | 5.254658 | 3.44 | Small CPxCG-related zinc finger protein; hypothetical protein |
| group_5199 | 0.039868 | 0.04512248 | 7.3125 | 2.98 | IS3 family transposase ORF A;transposase |
| group_5928 | 0.000958 | 0.03313556 | 12.04167 | 3.55 | Core-binding (CB) domain-containing protein;Shufflon-specific DNA recombinase;site-specific integrase |
| group_7021 | 0.001195 | 0.04975146<br>2 | 7.77443<br>6 | 2 | RAQPRD family<br>plasmid;Membrane<br>protein;Plasmid<br>Ignore RAQPRD<br>family |
| phzB | 0.001195 | 0.01239052 | 14.4677<br>4 | 3.56 | phenazine<br>biosynthesis protein<br>PhzB |
| phzF | 0.000122 | 0.01325153<br>3 | 9.64285<br>7 | 2.38 | Phenazine/pyocyanin<br>e biosynthesis<br>protein phzF;PhzF<br>family phenazine<br>biosynthesis<br>protein;phenazine<br>biosynthesis protein<br>PhzF |
| phzG~~~phzG<br>2 | 0.002828 | 0.04462022<br>2 | 5.6 | 1.87 | phenazine<br>biosynthesis FMN-dependent oxidase<br>PhzG;pyridoxine 5'-<br>phosphate oxidase C-terminal domain-containing<br>protein;pyridoxamin<br>e 5'-phosphate<br>oxidase |

### 3.8 Analysis of commonly identified gene families

We conducted a detailed literature review for each gene family that was commonly identified by both Scoary and PySEER.

#### 3.8.1 DsbA family protein

Proteins of the DsbA family play a key role in bacterial physiology by acting as thiol–disulfide oxidoreductases that catalyse disulfide bond formation in proteins transported to the periplasm of Gram-negative bacteria. This process is essential for correct protein folding, structural stability, and biological activity, particularly for proteins involved in virulence **(Xia et al., 2023).** Importantly, Furniss et al. demonstrated that disruption of DsbA-mediated disulfide bond formation severely compromises the activity of a wide range of β-lactamases **(Furniss et al., 2022).** Together, these findings suggest that DsbA proteins are critical for enabling antibiotic resistance enzymes to achieve their functional, active form.

#### 3.8.2 Copper resistance protein D

The copper resistance protein D (CopD) is a membrane-associated protein in Gram-negative bacteria that helps import essential copper from the periplasm into the cytoplasm, often working alongside the copper-binding protein CopC **(Lawton et al., 2016).** Such a system allows for the maintenance of the basal levels of copper necessary for the proper functioning of the cellular enzymes, even at high levels of copper stress **(Ladomersky & Petris, 2015).** Such mechanisms of copper homeostasis have been implicated in the development and maintenance of antimicrobial resistance in *P. aeruginosa*, implying that the stress response to metals indirectly contributes to the development of such resistance **(Virieux-Petit et al., 2022).**

#### 3.8.3 Transposase

Transposases are important enzymes in the spread of antibiotic resistance, which facilitate the movement of resistance genes within and between bacterial genomes **(Babakhani and Oloomi, 2018).** Transposases facilitate the movement of transposons, which insert the acquired genes into new genomic locations, sometimes creating complex mobile elements such as integrons and composite transposons. These elements enable bacteria to acquire multiple genes for resistance, thereby fueling the emergence of multidrug-resistant pathogens.

#### 3.8.4 Streptomycin 3’’-O-adenylyltransferase

The enzyme Streptomycin 3″-O-adenylyltransferase (AadA) confers resistance to streptomycin and spectinomycin by chemically modifying the antibiotics, which makes them unable to bind to the bacterial ribosome **(Papadovasilaki et al., 2015).** The carriage of the gene aadA in *P. aeruginosa* is generally the result of horizontal gene transfer, and this is frequently accompanied by the co-transfer of several resistance genes, which makes the development of multidrug resistance possible and allows the co-selecting of several resistance traits with the use of antibiotics **(Holbrook & Garneau-Tsodikova, 2018).**

#### 3.8.5 IS3 family transposase

IS3 family transposases play a role in antibiotic resistance in *P. aeruginosa* by facilitating genome rearrangements and gene disruption. This facilitates the integration of the IS3 family transposases into important genes, including the oprD porin gene, resulting in the inactivation of OprD and the reduction of imipenem uptake **(Bocharova et al., 2019).** Moreover, the activity of IS3 family transposases facilitates the spread of antibiotic resistance genes by mobile genetic elements.

#### 3.8.6 Small CPxCG-related zinc finger protein

The CPxCG-related zinc finger protein is also poorly characterised in *P. aeruginosa*, which is often annotated as a hypothetical protein by Panaroo. However, it has been reported that in Haloferax volcanii, this protein is involved in cell growth, swarming motility, biofilm formation, and has different metal-binding capabilities **(Üresin et al., 2024).** This suggests that this protein has functional significance, which is worthy of further exploration in *P. aeruginosa*.

#### 3.8.7 IS3 family transposase ORF A

In the case of *P. aeruginosa*, the IS3 family transposase ORF A is primarily a regulatory protein that regulates the expression of the active OrfAB transposase by programmed frameshifts **(Rousseau et al., 2004).** Although OrfA does not directly contribute to antibiotic resistance, it regulates mobile genetic elements that are involved in genome rearrangements. These elements are responsible for the horizontal spread of genes related to antibiotic resistance, such as β-lactamases and efflux.

#### 3.8.8 Shufflon-specific DNA recombinase

The shufflon-specific DNA recombinase, also called the Rci protein, is a site-specific enzyme that catalyzes DNA rearrangements in the shufflon region of bacterial plasmids **(Gyohda & Komano, 2000).** These rearrangements function as a genetic switch that introduces diversity in the structure of bacterial plasmids, which in turn helps in their spread by altering their specificity for bacterial recipients during conjugation.

#### 3.8.9 RAQPRD family plasmid

The RAQPRD family protein is commonly encoded within mobile genetic elements such as integrative conjugative elements and, in some cases, plasmids. Although its exact function remains unclear, it is thought to play a role in plasmid maintenance or conjugative transfer. Its consistent presence in genomic databases suggests it may contribute to the stability or spread of mobile genetic elements. **(InterPro.Ebi.ac.uk)**

#### 3.8.10 Phenazine biosynthesis protein PhzB/PhzF/PhzG

PhzB, PhzF, and PhzG are key enzymes in the phenazine biosynthesis pathway, where they act in a stepwise manner to convert chorismate into phenazine compounds such as phenazine-1-carboxylic acid (PCA) **(Pierson & Pierson, 2010).**

### 3.9 MLST Distribution and Association with ARGs and VFs

Using the seven standard housekeeping genes (*acsA, aroE, guaA, mutL, nuoD, ppsA, and trpE*), we attempted to assign sequence types (STs) to all 95 *P. aeruginosa* genomes. The MLST analysis successfully identified STs for 70 out of 95 samples, while the remaining genomes could not be assigned to a known ST. The exact details of the ST assignments for each isolate are given in Supplementary File 6.

As indicated in **Figure 7A**, the different sequence types varied in their occurrence in the data set. Among the different sequence types, the highest occurrence was of sequence type 233. It is worth pointing out that all the isolates of sequence type 233 were imipenem-resistant. This is in line with the literature indicating the international high-risk clone of sequence type 233, which is associated with extensive drug resistance (XDR) and severe hospital-acquired (nosocomial) infections **(Del Barrio-Tofiño et al., 2020).**

Another sequence type that was found to be frequent in the study is sequence type 235. This sequence type included imipenem-resistant and imipenem-susceptible isolates. Sequence type 235 is also well recognized as a globally prevalent high-risk clone **(Del Barrio-Tofiño et al., 2020).** Besides the two sequence types, sequence type 389 was found to be present in a large number of isolates.

**Figure 7B** shows the relationship between resistance and virulence at the sequence type level. This is indicated by the mean AMR gene count against the mean virulence factor count for each of the sequence types found.

Among the most distributed sequence types, ST 233 showed the highest combined mean values of AMR and VF genes, highlighting its strong association with both resistance and virulence. ST 235 also exhibited elevated values, although lower than those observed for ST 233.

Interestingly, ST 964 showed a high richness of both AMR and VF genes. However, as seen in **Figure 7A**, this ST was present in only a small fraction of the total genomes, indicating that while it is gene-rich, it is less prevalent in the dataset.

**Figure 7:**
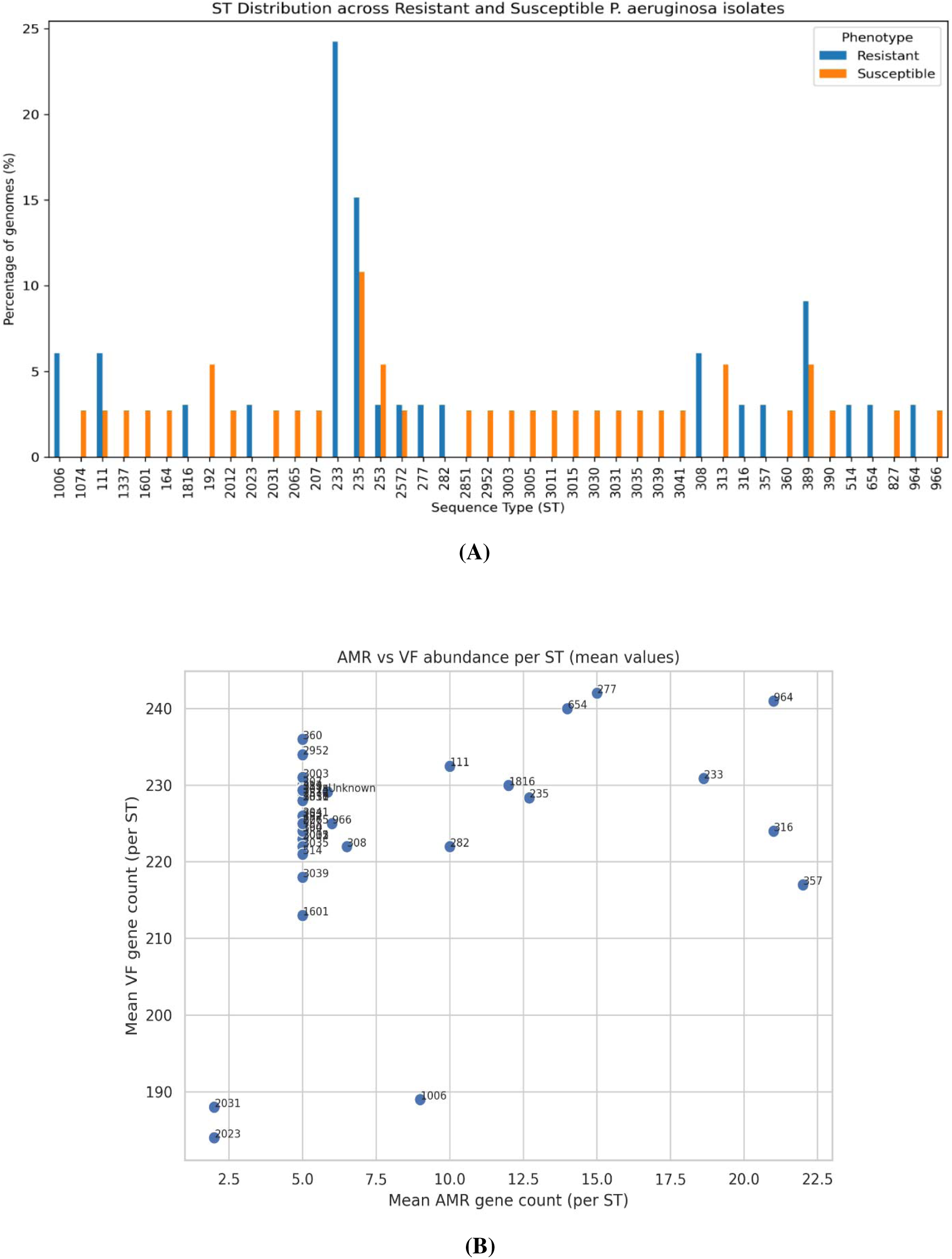
Multilocus sequence typing (MLST) analysis of imipenem-resistant and imipenem-susceptible *P. aeruginosa* isolates. **Figure (A)** shows the richness of the identified sequence types (STs) across all strains. **Figure (B)** illustrates the average number of antimicrobial resistance (AMR) genes and virulence factor (VF) genes per ST. *Both the figures were generated using Python-based libraries*.

### 3.10 Machine Learning Model Performance and Validation

We developed a LASSO logistic regression model to predict imipenem resistance in *P. aeruginosa* using twelve genes identified from GWAS analysis. To obtain an unbiased estimate of model performance while accounting for clonal relatedness, leave-one-sequence-type-out cross-validation was applied. This strict validation approach ensured that the model was evaluated on genetically distinct lineages rather than closely related strains. The model showed strong predictive ability with an AUC-ROC of 0.836 (**Figure 8A**), indicating good discrimination between resistant and susceptible isolates. Performance variability was also observed for each type of sequence, primarily due to the sample size and class distribution; however, the results show the ability of the model to generalize over unseen sequence types.

#### 3.10.1 Feature Importance and Biological Interpretation

The LASSO analysis also identified a subset of genes associated with GWAS, which contributes to the prediction of resistance after embedded feature selection. As presented in **Figure 8B**, most genes were associated with non-zero coefficients, implying their contribution to the predictive model, while others were associated with negative or minimal coefficients, implying weaker or negative associations after accounting for the correlations between features. Of all the features, the highest positive value was associated with group_7021, followed by phzB and phzF, implying that these are the most important features in the model. Moderate positive values were associated with group_4007, group_4947, group_10894, group_5928, group_10887, and group_3484, implying their contribution to the model. However, negative values were associated with group_10880, phzG-phzG2, and group_5199, implying that the presence of these features is associated with a lower probability of resistance.

In terms of functional significance, many of the predictors highlighted in the model are associated with mobile genetic elements, such as transposases and integrases, which suggests the importance of the accessory genome in the resistant strains. The contribution of the genes associated with phenazine biosynthesis also supports the idea that metabolic and redox adaptation may be important in supporting the survival of the cells under antibiotic stress. Overall, the LASSO model builds on the GWAS results and highlights a small number of genes that work together to support the predictive model for imipenem resistance in *P. aeruginosa*.

### 3.11 Confusion Matrix and Per-Gene Performance

From the confusion matrix (**Figure 8C**), it is clear that the LASSO logistic regression model was performing reliably in differentiating between imipenem-resistant isolates and susceptible isolates. A larger number of susceptible isolates were correctly identified (42 true negatives), although there were also instances of susceptible isolates being incorrectly identified as resistant (12 false positives). On the other hand, it was also clear that the majority of resistant isolates were correctly identified (32 true positives), although there were instances of isolates being incorrectly identified as susceptible (9 false negatives). This is not surprising, considering the diverse nature of the dataset and the fact that the LASSO logistic regression model is based on gene presence-absence patterns alone. These results clearly indicate that the LASSO logistic regression model is maintaining an appropriate balance between being sensitive and specific, although it is slightly more sensitive when it comes to susceptible isolates.

The analysis of performance per gene (**Figure 8D**) also indicates that there is variability in predictive performance for each gene. For example, group_5199, group_4007, and group_5928 had relatively lower false positive contributions, suggesting better individual specificity for these genes. In contrast, phzB and the phzG–phzG2 locus showed higher false-positive signals, suggesting that although these genes are associated with resistance, their presence alone is not sufficiently discriminative. Together, these findings support the use of the multivariate LASSO framework, which integrates information across multiple loci and therefore provides a more robust prediction of imipenem resistance in *P. aeruginosa* than any single gene marker alone.

**Figure 8.**
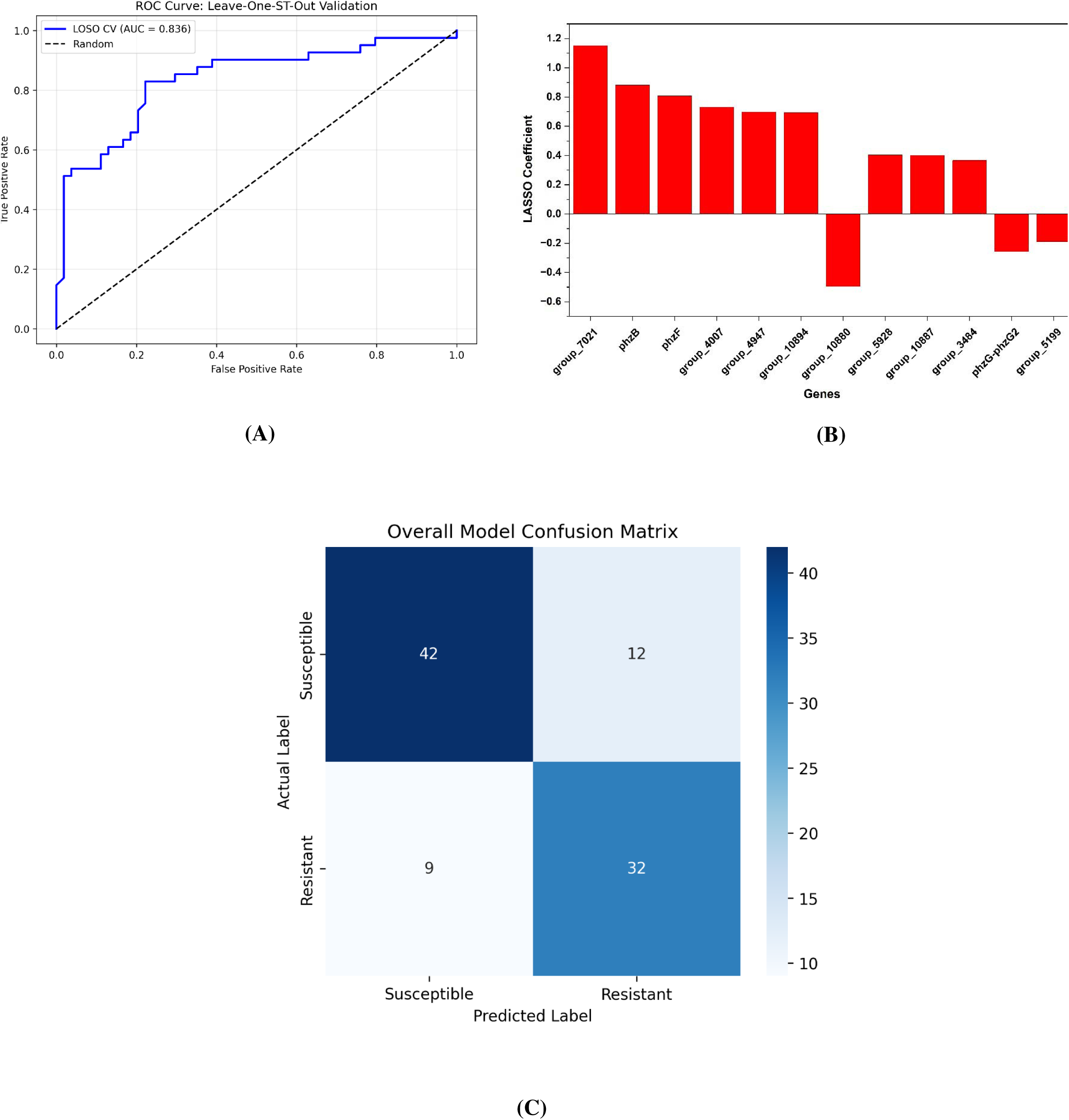
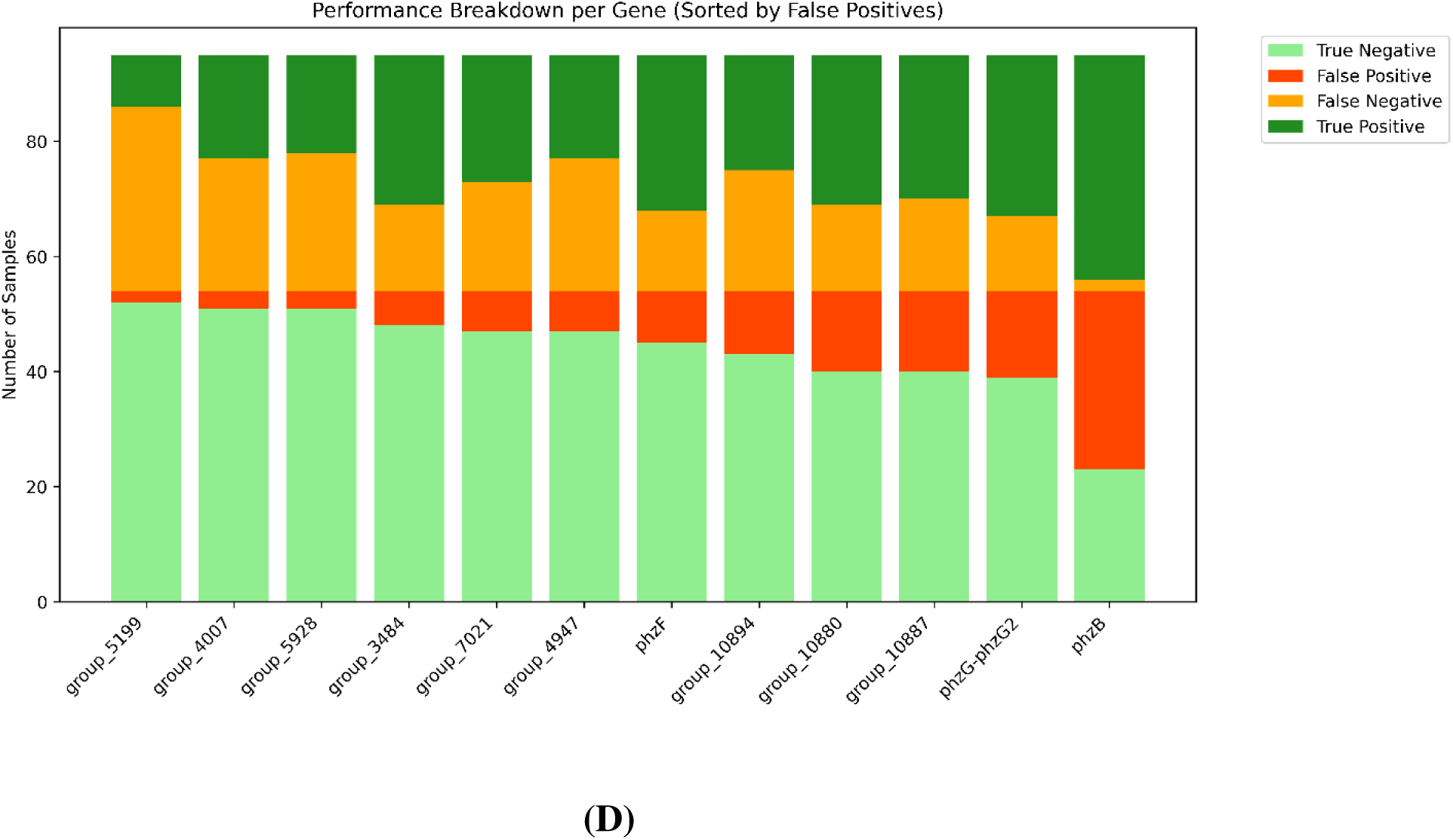
Multivariate and univariate prediction of imipenem resistance. **Figure (A)** shows Receiver operating characteristic (ROC) curve obtained using leave-one-sequence-type-out (LOSO) validation. **Figure (B)** illustrates LASSO coefficient profile highlighting features contributing to resistance prediction. **Figure (C)** shows the confusion matrix summarizing classification performance of the LASSO logistic regression model. **Figure (D)** shows Per-gene diagnostic performance evaluated using the univariate presence–absence approach. *Figure A, C & D were generated using Python-based libraries and Figure B was generated using Origin 2025b (*https://www.originlab.com/)

### 3.12 Resistant-Specific Gene Co-occurrence Network

A resistant-specific gene co-occurrence network was constructed using the filtered pan-genome matrix together with phenotype metadata (**Figure 9**). After applying the correlation threshold (|ρ| ≥ 0.3) and FDR correction, a tightly connected accessory gene module emerged. Ten highly connected nodes were found using the degree centrality analysis, as depicted in **Figure 9**. These nodes were classified as hub genes (**Table 3**), where each of the hub genes exhibited perfect correlation values (|ρ| = 1).

Hub genes were ranked according to their degree centrality values. The top ten hub genes were selected to cover the most important nodes without incorporating less connected genes, which might have introduced “noise” in the analysis. This selection criterion offered a fair balance of the accessory genes related to resistant *P. aeruginosa*.

**Table 3:** Top ten hub genes identified from the resistant-specific gene co-occurrence network, along with their functional annotations.

| Gene | Annotation |
| --- | --- |
| sfnB | SfnB family sulfur acquisition oxidoreductase |
| group_1341 | Luciferase-like domain-containing protein |
| group_5873 | cell division protein ZapE |
| group_9305 | HTH lysR-type domain-containing protein |
| msuE | FMN reductase |
| group_1791 | ABC transporter substrate-binding protein |
| tssA | type VI secretion system protein TssA |
| group_10685 | Phage tail protein |
| group_10665 | hypothetical protein |
| group_4235 | Acyl-CoA dehydrogenase protein; Monooxygenase |

Functionally, the hub set suggests that there are coordinated changes in metabolic capability, cellular fitness, and accessory genome features in resistant populations. Some of the genes in the set are related to redox reactions and energy metabolism, implying that resistant populations might be in a heightened metabolic state. Moreover, the presence of a type VI secretion system component suggests that there might be a role for inter-bacterial competition in resistant populations. In addition, the presence of regulatory and transport-related genes suggests that there might be a coordinated effect in the genome associated with resistance. It should also be noted that, with the identification of phage-related and hypothetical proteins, we are probably witnessing the implication of poorly understood accessory components, thus supporting the suggestion that horizontal transfer of genetic material and lineage-specific genomic content are important in explaining the observed resistant population structure.

It should also be stressed that these hub genes should not be taken as causal agents of imipenem resistance. They are, as expected, markers of resistant genomic background.

**Figure 9:**
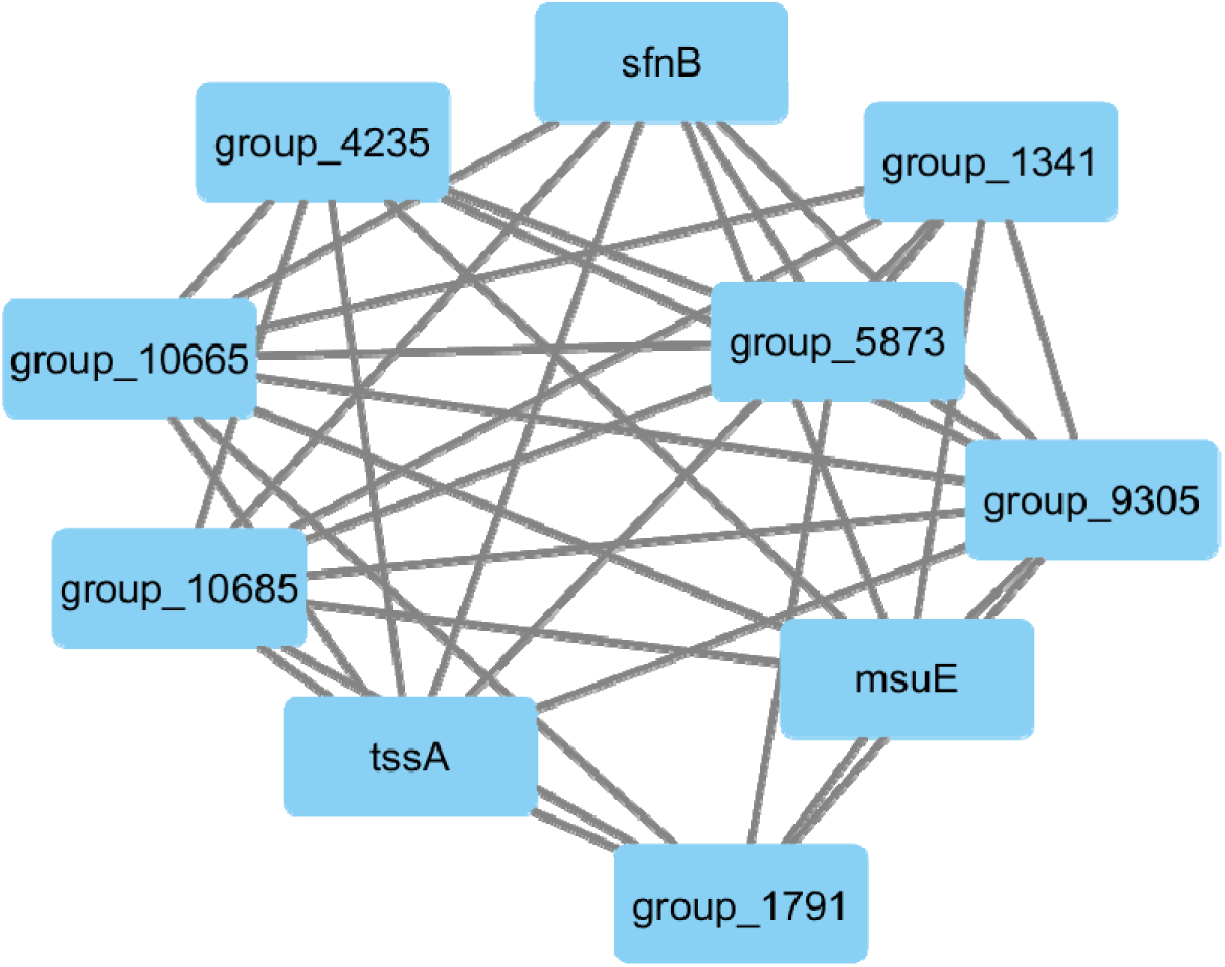
Visualization of the resistant-specific gene co-occurrence network highlighting the top ten hub genes identified based on degree centrality. *Figure was generated using Cytoscape_v3.10.3 **(Shannon et al., 2003)***

### 3.13 Weighted Gene Co-expression Network Analysis (WGCNA)

#### 3.13.1 Soft-Thresholding Power Selection

For the construction of the weighted gene co-occurrence network based on the pangenome presence/absence data, the soft-thresholding power value was calculated based on the scale-free topology criterion. The range of the soft-thresholding power value was set from 1 to 20. Based on the scale-free topology criterion, as depicted in **Figure 10A & 10B**, the values of the signed R² did not follow a linear trend; the values also did not reach the 0.90 benchmark. The highest value was recorded at a soft-thresholding power value of 6, beyond which the curve plateaued. Such a behavior is expected in binary pangenome data, which do not necessarily exhibit the same distributional characteristics as continuous data in the form of transcriptomic data. Moreover, the assessment of the mean connectivity profile also provided support for this choice, as at a low power level (β=1), the network was found to be both dense and undifferentiated. However, as the power level increased, the weak and potentially noisy connections were reduced, and at a power level of 6, the network had transformed to a more differentiated state with much lower mean connectivity values. According to WGCNA guidelines, when the desired scale-free fit cannot be achieved, the lowest power that provides a reasonable approximation to scale-free behavior while maintaining sufficient connectivity should be selected (**Zhang & Horvath, 2005**). Therefore, β = 6 was chosen for downstream analysis as it provided the best balance between network sparsity and preservation of biologically meaningful signal in the present pangenome dataset.

#### 3.13.2 Pangenome Network Construction and Module Identification

We constructed a co-occurrence network from the pangenome presence-absence matrix to identify clusters of genes that share similar distribution patterns across the isolates. A hierarchical clustering tree was generated based on the topological overlap matrix, and the dynamic tree-cut algorithm was used to partition the genes into distinct modules. In order to make our network robust, we also merged modules that had very similar eigengene profiles, with a dissimilarity threshold of 0.25 as indicated in Supplementary File 7. This refinement step gave us a consolidated set of modules that each represented a unique pattern of gene co-occurrence in the pangenome.

#### 3.13.3 Analysis of Module-Trait Relationships

After the identification of the modules, the correlation analysis has been conducted to associate the gene clusters with the resistance phenotype. This has helped us to identify the specific regions of the genome that are significantly associated with the phenotypes of interest. Total 154 different modules have been identified. All the statistical data such as correlation values, gene numbers present in each module, and other statistical data are provided in Supplementary File 8-9. From the total, the top 10 modules that show high correlation with the resistance phenotype, as depicted in **Figure 10D**.

The strongest correlation was observed with the black module, with a correlation coefficient of r = 0.50, p < 0.001, showing a moderate but significant relationship with resistance. Next in rank was the royalblue3 module, with a correlation coefficient of r = 0.43, p < 0.001, followed by the magenta and grey modules, with a correlation coefficient of about r = 0.42, p < 0.001. Other modules, including blue, lightcyan, grey60, plum4, white, and mediumpurple2, also showed significant positive correlations with resistance.

As the model of module-trait analysis revealed, the polygenic nature of the resistance phenotype is underscored. Although the black module had the highest correlation, the magnitude was also moderate at 0.50. This suggests that a single module does not account for the entire phenomenon of resistance, as several modules contributed equally to the signal, supporting the theory that the data are multifactorial in nature.

Notably, the moderate correlation values are also expected for the presence/absence pangenome data, as binary data generally have weaker network associations than expression data. However, the consistent positive associations for the top modules suggest that the WGCNA method correctly captured the biological co-occurrence patterns that are relevant for resistance. These resistance-associated modules provide a focused set of candidate gene clusters for downstream hub gene identification and functional characterization.

**Figure 10:**
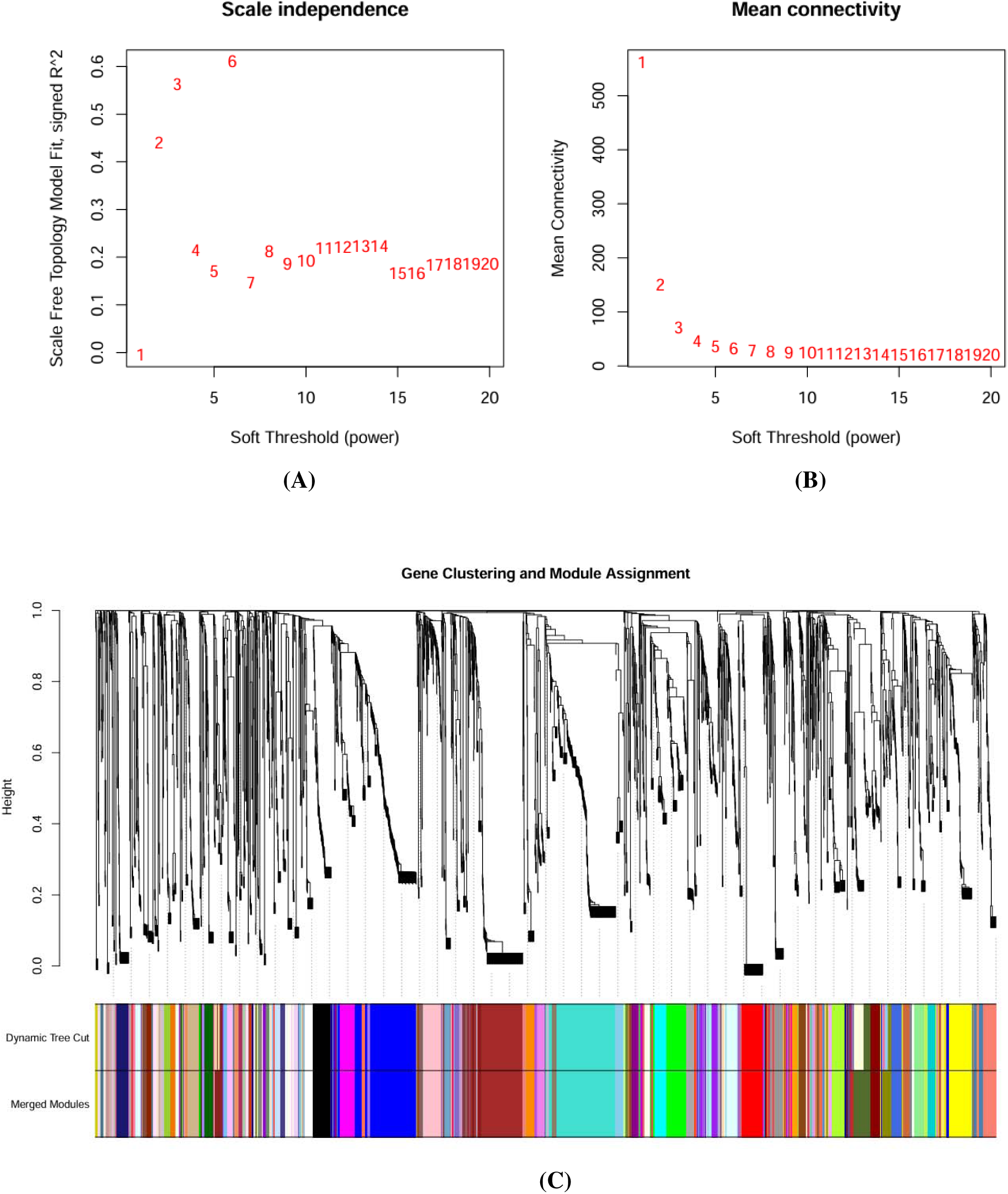
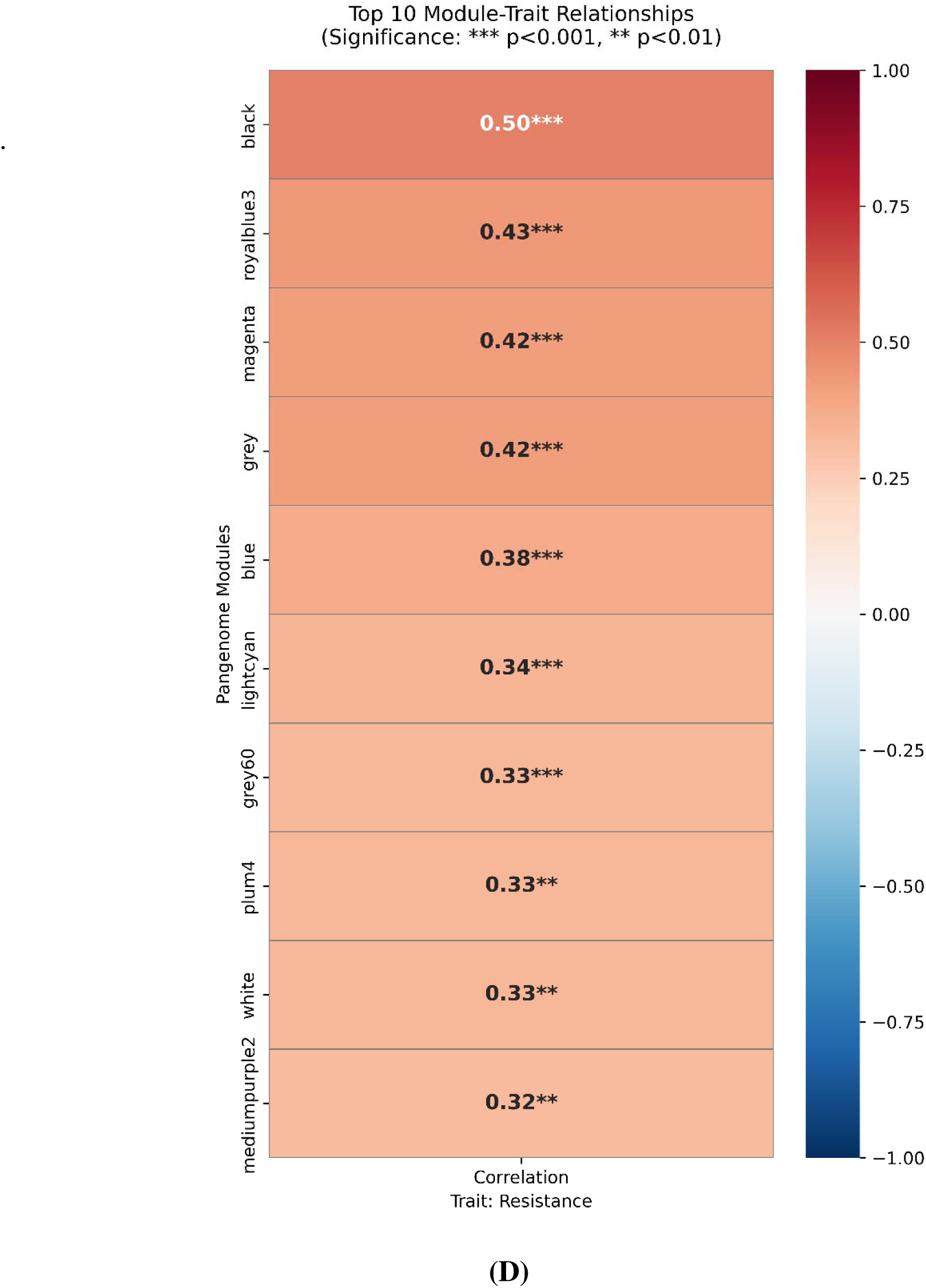
WGCNA soft-threshold selection, module detection, and module–trait relationships **Figure (A)** shows scale-free topology fit index (scale dependence) across candidate soft-threshold powers used to determine the appropriate network construction parameter. **Figure (B)** shows mean connectivity corresponding to the tested soft-threshold powers, illustrating the trade-off between scale-free topology and network connectivity. **Figure (C)** illustrates Hierarchical clustering dendrogram of genes with dynamic tree cut–based module assignment, showing the identified co-expression modules. **Figure (D)** shows heatmap of the top ten module–trait relationships, highlighting modules most strongly associated with the resistance phenotype. *Figure A, B & C were generated with the help of R studio software and Figure D was generated with the help of Python libraries*.

#### 3.13.4 Identification of Key Hub Genes in Resistance Modules

To identify the most influential genes within the resistance-associated modules, we calculated “hub scores” by multiplying module membership (the correlation of a gene with its module) by gene-trait significance (the correlation of a gene with the resistance phenotype). This dual-filtering approach ensures that the identified hub genes are not only central to the network’s structure but also highly relevant to the biological trait. A complete list of hub genes across all modules, along with their statistical details, is provided in Supplementary File 10. Among the modules, the black module exhibited the strongest association with the resistance phenotype. The key gene products in this module, according to their order in **Table S1** of Supplementary File 11, encompass DNA binding and maintenance, mobile genetic elements, transcriptional regulation, transport, and other gene products with uncharacterized domains. The significance of functions including integrase, partitioning proteins, and transport activities indicates that genome plasticity, transcriptional control, and transport activities could be involved in conferring resistance.

#### 3.13.5 Integration of WGCNA Hub Genes with GWAS Findings

To further validate our biological confidence in our network results, we also compared our WGCNA hub genes with previously identified candidate genes from the Genome-Wide Association Study (GWAS). Among the top 12 GWAS-identified genes, four of them are also hub genes in our WGCNA results, including group_10880 and group_10887, which are located in the darkred module, as well as group_4947 and phzB, which are hub genes in the plum3 and grey modules, respectively. GWAS results are based on statistical associations between genetic loci, whereas WGCNA results are based on coordinated gene behaviour. Thus, these results provide strong biological confidence that these genes are indeed associated with resistance.

We also went on to investigate which other hub genes these validated candidates interact with in their respective modules. In the darkred-colored module, group_10880 and group_10887 appear to be part of a conjugative transfer machinery, connecting with seven other hubs including TraG family proteins, a relaxase, and multiple integrating conjugative element proteins as listed in **Table S2** of Supplementary File 11. The fact that there is also a DNA repair protein included in this module indicates that it may be involved in coordinating the movement of genetic elements as well as DNA damage response for resistance.

Interestingly in the plum3 module, the group_4947 cluster is found together with genes related to secondary metabolic and genomic rearrangement processes, such as a saccharopine dehydrogenase, transposases, integrases, and uncharacterized proteins. The TcmP family protein and the cointegrate resolution protein also indicate the occurrence of active DNA recombination events in this module. This module indicates a mixed contribution of genome mobility and cell adaptation processes.

Lastly, phzB in the grey module is at the center of phenazine synthesis, directly interacting with phzA and surrounded by redox proteins such as rubredoxins and an acetyltransferase.

Transcriptional regulators and translation machinery are also part of the complex, implying that phzB-mediated resistance is achieved by the synthesis of antimicrobial agents as well as metabolic regulation.

### 3.14 Network Topology and Degree Distribution

In order to study the global architecture of the resistant and susceptible gene co-occurrence networks, the node degree distribution of the networks was compared (**Figure 11A**). It is found that the resistant network has a right-shifted distribution compared to the susceptible network. This suggests that the genes in the resistant network have a greater number of connections compared to the susceptible network. This is also reflected from the fact that the average node degree of the resistant network (153.7) is significantly higher compared to the susceptible network (115.9).

The degree distribution of both networks had a long-tailed distribution on the log-scaled y-axis, implying the presence of highly connected nodes as well as nodes with fewer interactions. The resistant network had a wider distribution towards higher degrees, implying a more dense and connected network in the resistant strains.

### 3.15 Global Network Topology Remains Largely Conserved

The global topology analysis showed that the gene co-occurrence networks of resistant and susceptible isolates have similar global topological features, as demonstrated in **Figure 11B**. Although the resistant isolate’s gene co-occurrence network had a slightly higher density, clustering coefficient, and transitivity compared to the susceptible isolate’s gene co-occurrence network, the differences were very small. This result suggests that the global architecture of the pangenome network is similar in both phenotypes. This leads us to conclude that imipenem resistance in this dataset is very unlikely to be caused by major changes in the network architecture.

### 3.16 Module-Level Connectivity

Significant differences in the pattern of connectivity within modules between resistant and susceptible strains were revealed by the comparison of the intramodular connectivity, as depicted in **Figure 11C**. Of the six modules found to be highly connected within the resistant network, most of the modules showed significantly increased connectivity within the resistant strains, suggesting increased co-occurrence of genes within the resistant phenotype. The turquoise module showed the greatest differential connectivity, showing increased connectivity within the resistant compared to the susceptible phenotype. A similar pattern, although more dramatic, was observed for the brown module, where connectivity was high within the resistant phenotype and almost absent within the susceptible phenotype.

In contrast, the red module had a similarity in the values of connectivity between the two phenotypes, implying that the intrinsic structure of this module is likely to be stable and consistent, irrespective of the resistance state. The blue module had a sharp decrease in the values of connectivity in the susceptible strains, while the yellow and black modules had a decrease in the values of connectivity in the susceptible strains, although this decrease was moderate.

Overall, the above results suggest that modules highly organized in resistant strains do not necessarily maintain the same level of internal connectivity in susceptible strains. Instead, it appears that there is a relationship between resistance and the selective strengthening of certain gene co-occurrence modules.

#### 3.16.1 Module Connectivity Patterns of GWAS–WGCNA Overlapping Modules

We then went on to examine the connectivity pattern for the three modules that had shared genes between GWAS and WGCNA analysis results. A phenotype-dependent pattern was evident, as demonstrated in **Figure 11D**, although the extent of change varied between modules.

In the darkred module, the mean intramodular connectivity in the susceptible strains was found to be higher than in the resistant strains, which suggests that the co-occurrence of genes in this module somehow becomes weaker in the resistant conditions. A similar trend, although not as obvious as in the darkred module, was also found in the plum3 module, in which the susceptible strains had a slightly higher connectivity than the resistant strains.

On the contrary, the grey module had very low connectivity for both phenotypes, consistent with the typical interpretation of the grey module in WGCNA as a set of weakly or unassigned genes. The similarity between the resistant and susceptible groups for the grey module also suggests that there was limited co-occurrence for its genes. Collectively, the module results suggest that resistance is associated with changes in the organization of the networks within relevant biological gene sets, as supported by both GWAS and WGCNA data.

**Figure 11:**
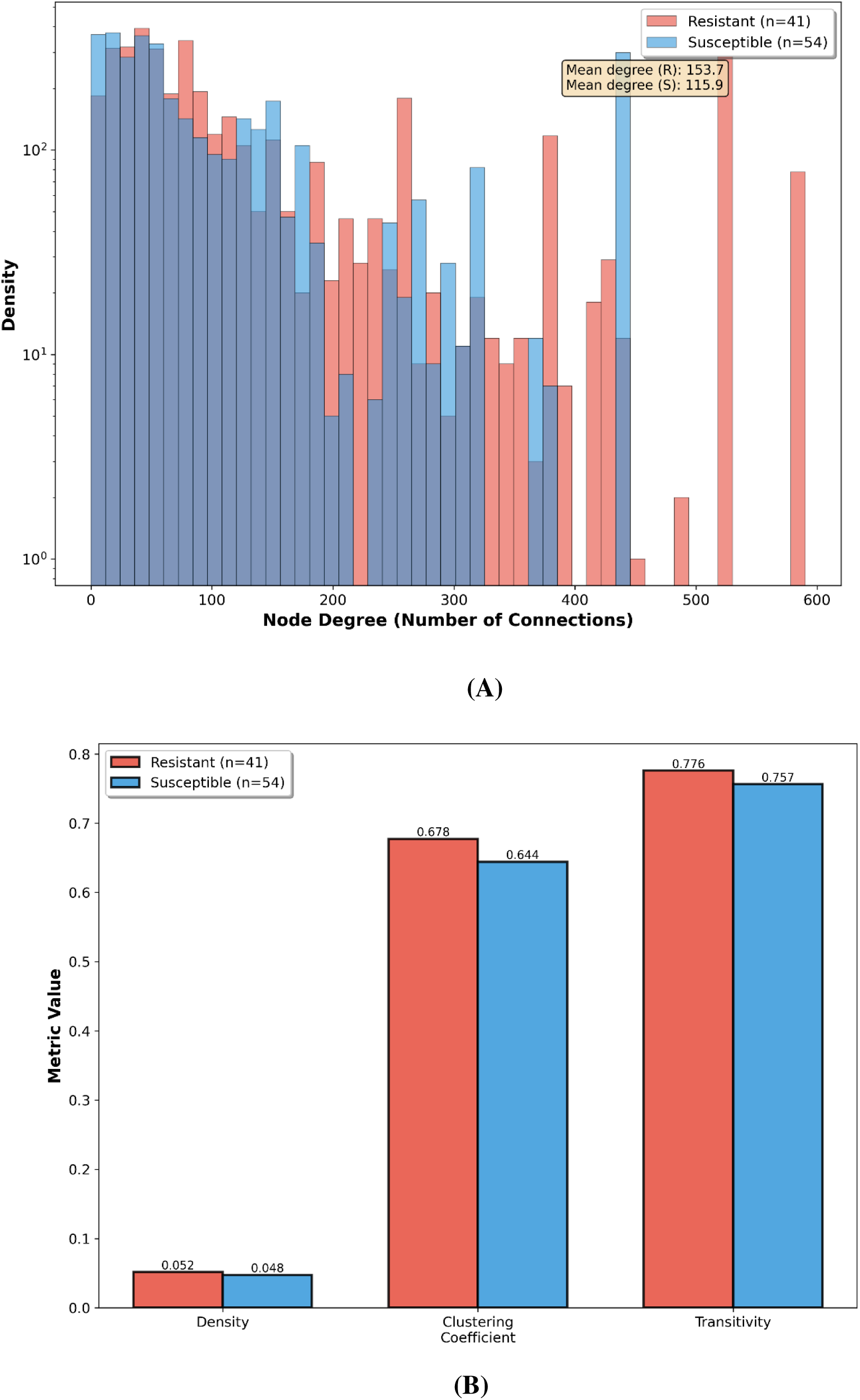
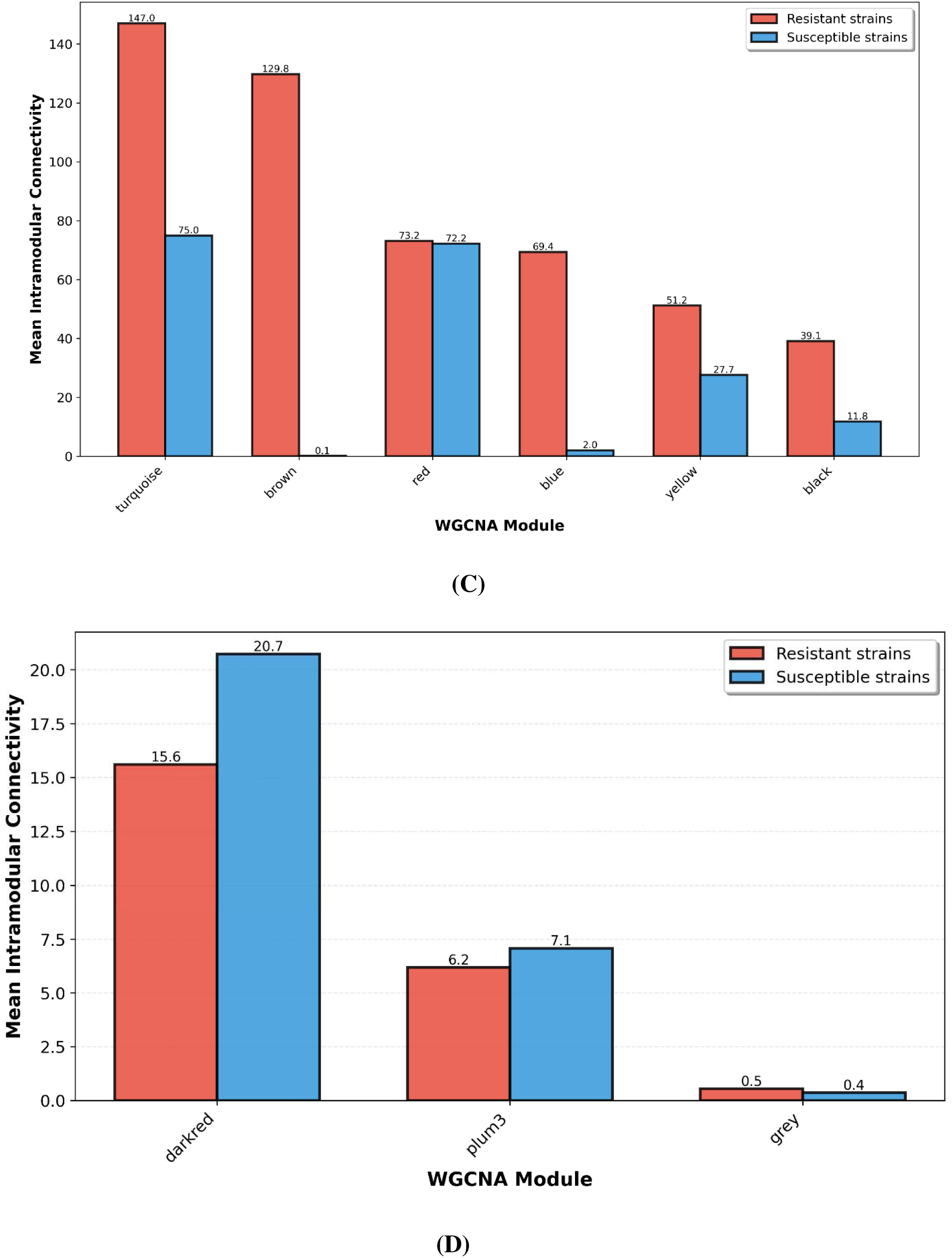
Comparative network analysis of resistant and susceptible *P. aeruginosa* strains. **(A)** Degree distribution of gene co-occurrence networks for resistant and susceptible groups. **(B)** Global network topology metrics comparing the two phenotypes. **(C)** Mean intramodular connectivity (kWithin) across the top six modules in resistant versus susceptible strains. **(D)** Targeted comparison of mean intramodular connectivity for the darkred, plum3, and grey modules containing GWAS–WGCNA overlapping genes. *All the figures were generated with the help of Python libraries*.

### 3.17 Silent Resistome Activation Model: A Novel Hypothesis

The Silent Resistome Activation Model (SRAM), which is based on genomic, statistical, and network analyses of data, describes the formation of resistance to imipenem in *P. aeruginosa.* The key postulate of this theory is that the development of resistance occurs not via a simple acquisition of genes but through the stepwise activation of a pre-existent silent resistome.

Through our genome-wide association analysis, we have discovered that imipenem resistance is inherently polygenic, which means that it involves multiple resistance genes; namely, twelve genes instead of one. Compared to the susceptible isolates, the resistant isolates exhibit a wider range of antibiotic resistance genes. In susceptible isolates, carbapenemase-producing blaOXA genes have been found, although they are inactive because they lack insertion sequences for activation and weak regulatory sites. More importantly, the lack of DsbA-like oxidoreductases and related proteins limits the appropriate development of the enzyme, thus OXA cannot form catalytic enzymes.

Activation of resistance is accomplished through extensive modification of the accessory genome using mobilizable genetic elements like transposases, integrases, and plasmid DNA. They contribute to increased dosage of genes, provide strong promoters, and secure resistance loci. Weighted gene co-expression network analysis reveals that the process entails selective reorganization of certain gene neighborhoods and not an entire network change. The darkred module, which is associated with conjugation transfer proteins, demonstrates increased connectivity among sensitive strains, implying that resistance is caused by partial reconfiguration of the network. In contrast, the plum3 module has been shown to be involved in genome rearrangements and secondary metabolites, implying resistance evolves through genome plasticity.

The next interesting aspect is the metabolism and redox changes. Genes responsible for phenazine biosynthesis and especially phzB were among those constantly featured as the most significant factors in resistance regardless of the fact that they are not classical resistance factors. Being a key factor located around redox and regulation elements, it seems plausible to assume that phzB creates favorable redox state in periplasm needed for OXA fold and activity, thus turning latent resistance potential into an enzyme production. Certain genes showing near-perfect resistance specificity in univariate analysis yet negative coefficients in multivariate models highlight the importance of population-aware approaches for interpreting accessory genome signals.

Evolving from an evolutionary perspective, the preponderance of sequence type 233 among resistant strains designates it as a dangerous strain, although poor phylogenetic clustering suggests separate acquisitions of resistance within diverse genetic backgrounds using accessory genomic mechanisms. The robust predictive power of our LASSO algorithm (AUC = 0.836) confirms that resistance prediction must be assessed based on both genome background and cellular environment. Thus, *bla*OXA functions as a conditional resistance element dependent upon supporting genomic architecture including specific mobile elements, redox-processing capabilities, and metabolic adaptation systems.

The proposed theory is not restricted to *bla*OXA and *P. aeruginosa* but could be applicable to a broad range of antibiotics and bacteria, underscoring the necessity that surveillance programs track the activation capacity of the silent resistome rather than just tracking known resistance. Verification of the phenazine biosynthetic pathway enzymes and the DsbA family of oxidoreductases via transcriptional, metabolic, and temporal studies will validate these connections. The model is depicted in **Figure 12**.

**Figure 12:**
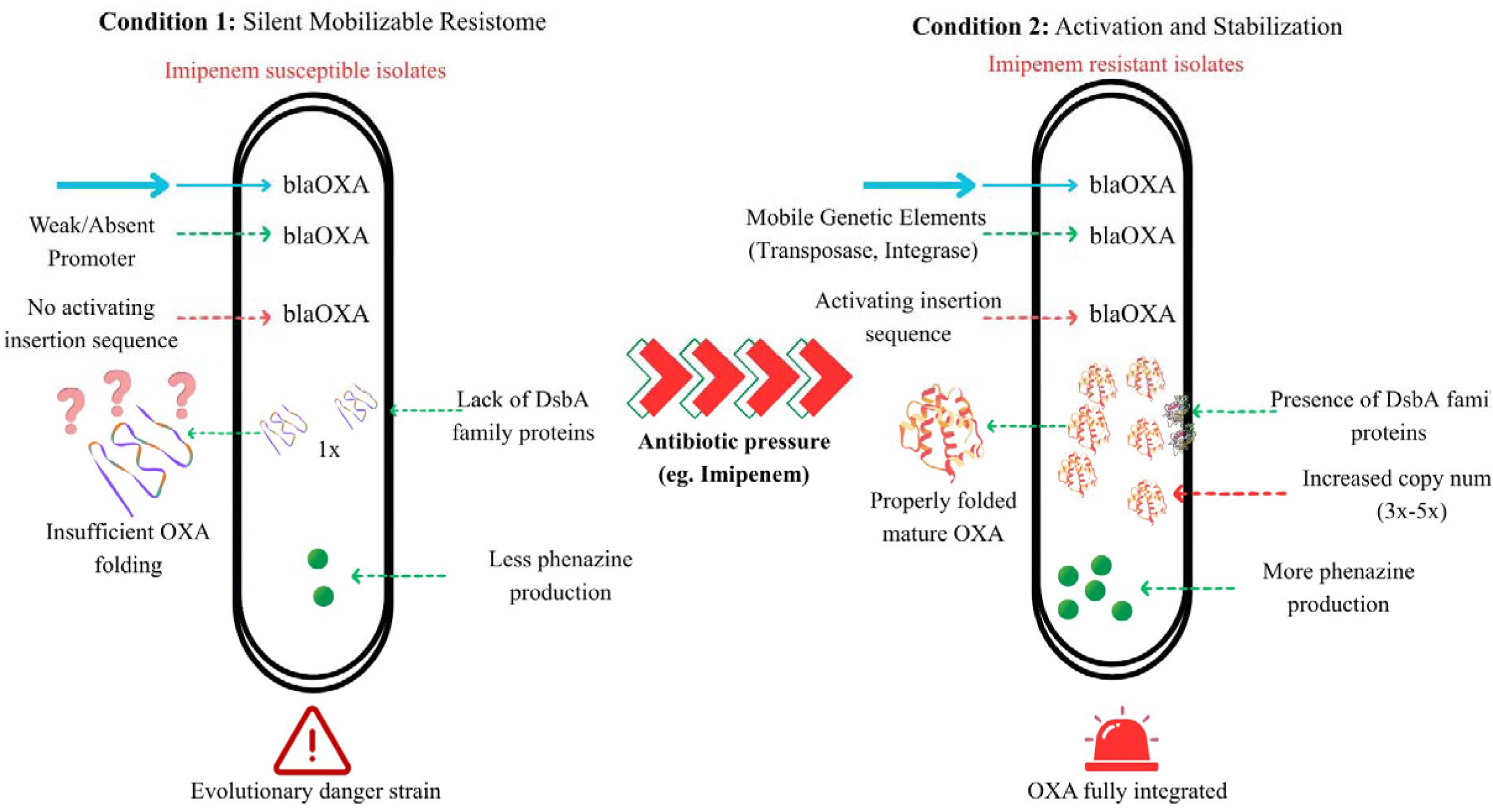
A summary schematic illustrating the proposed novel hypothesis, the “Silent Resistome Activation Model,” developed based on the overall findings of this study. *The figure was created using an online designing tool, Canva* (https://www.canva.com/).

## Conclusion

The present work demonstrates that resistance of *P. aeruginosa* to imipenem is not a consequence of a specific gene but rather the result of a coordinated effort of several genes acting in concert. The current research has demonstrated that there is a number of genes associated with resistance, such as blaOXA, even in susceptible strains, which can be inactive under certain conditions unless they receive proper genomic and cellular context support. On the basis of the same, we propose the Silent Resistome Activation Model, where resistance develops through the activation of hidden genetic potential. These findings majorly highlight the need to tap into the reservoir of silent genes for genomic surveillance and prevention of AMR. These findings also highlight the need to move beyond single-gene diagnostics and focus on genome-wide approaches to better predict and control antibiotic resistance.

## Author Contributions

Sarfraz Anwar and Sintu Kumar Samanta contributed to the study conception and design. Material preparation, data collection, and analysis were performed by Sarfraz Anwar, A R Aromal and Ananya Anurag Ananad. The first draft of the manuscript was written by Sarfraz Anwar. Sintu Kumar Samanta and Ananya Anurag Ananad edited the manuscript and provided advice on the experiments. All authors commented on previous versions of the manuscript. All authors read and approved the final manuscript.

## Acknowledgments

Sarfraz Anwar would like to thank University Grants Commission (UGC) for his fellowship. Ananya Anurag Anand and A R Aromal would like to thank MoE, Govt. of India for their fellowship. We are thankful to IIITA for providing research facility.

## Statements and Declarations

### Ethical approval

Not applicable

### Informed consent

Not applicable.

### Clinical trial number

Not applicable.

