## Supplementary figures and images for "Genome-Wide in silico analysis reveals activation of a silent resistome driving imipenem resistance in *Pseudomonas aeruginosa*"

### Supplementary File 3

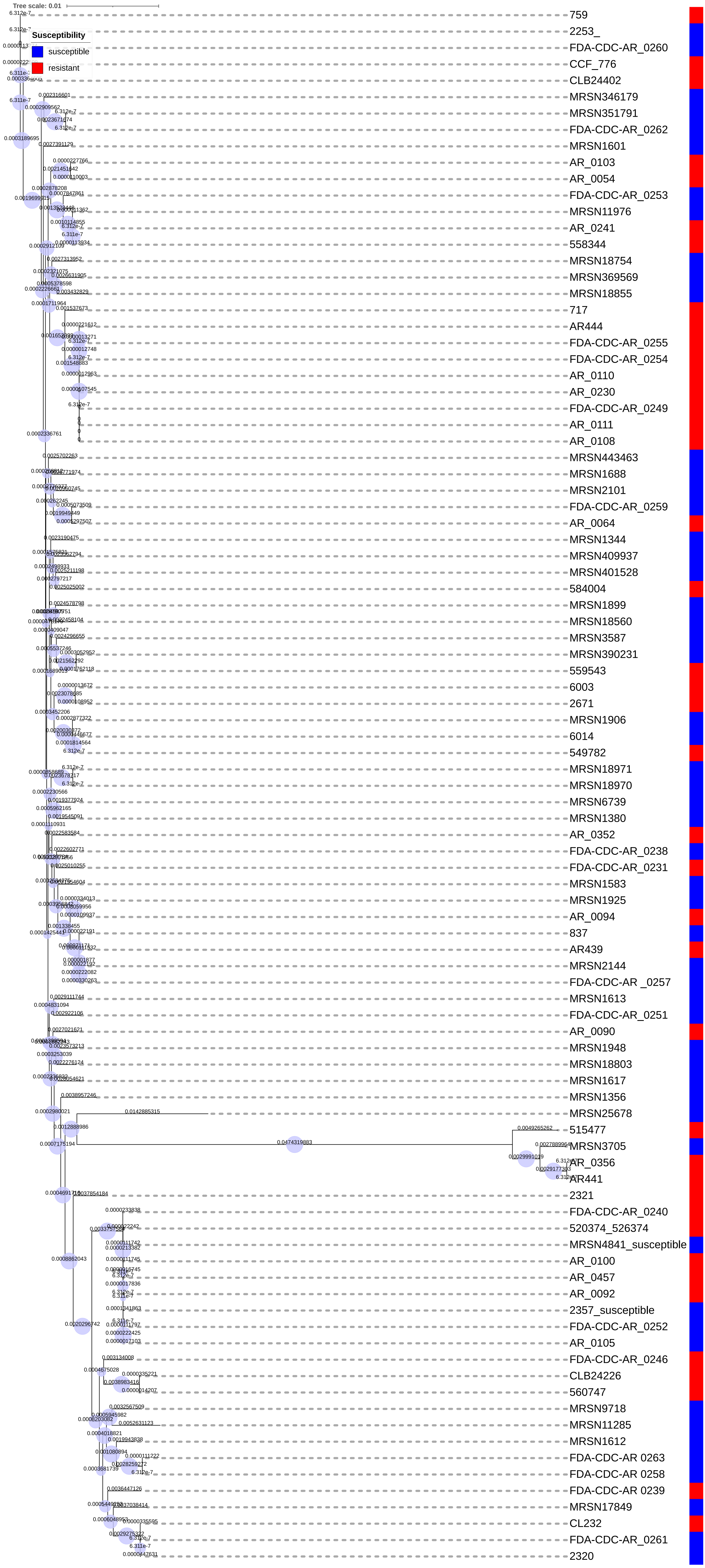

### Supplementary File 7

#### Clustering of module eigengenes

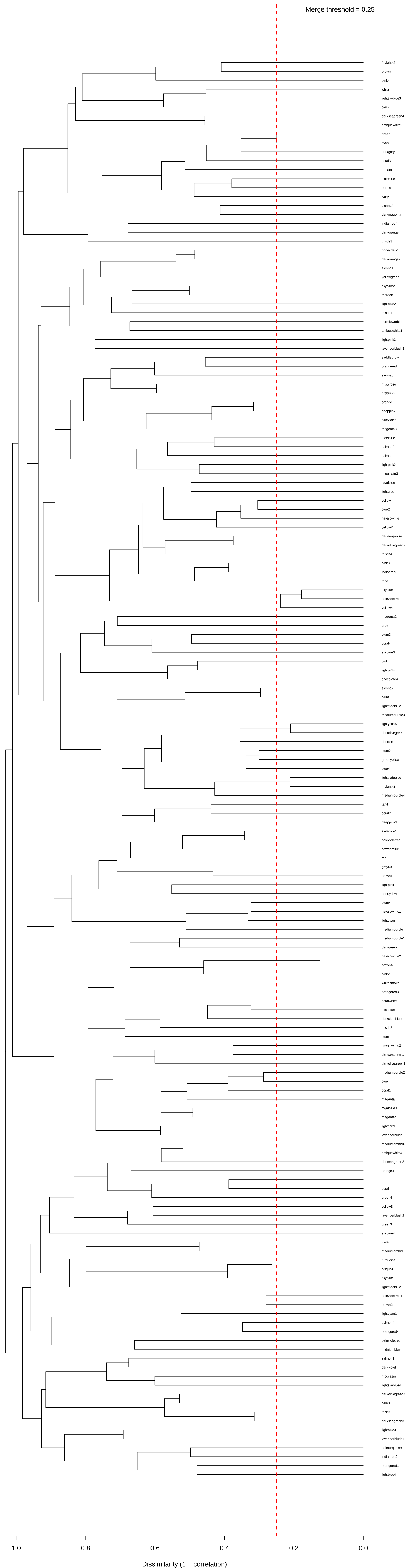

### Supplementary File 8

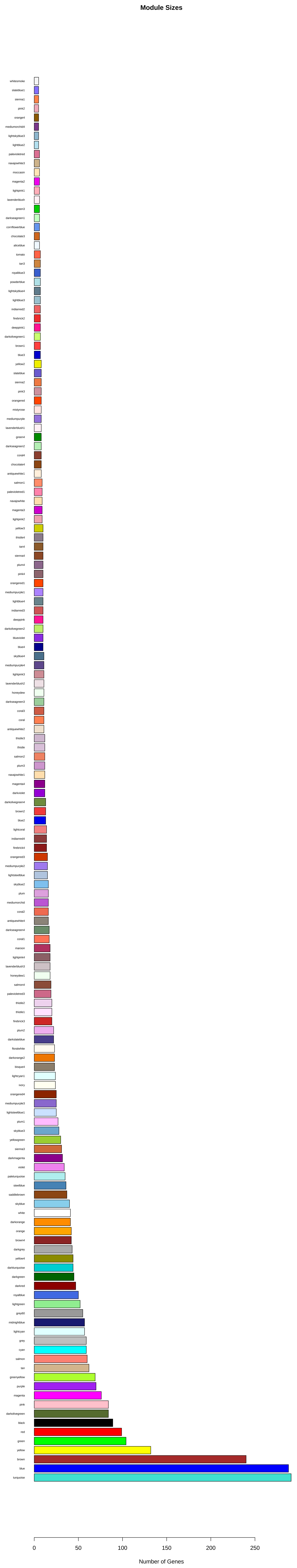

### Supplementary File 9

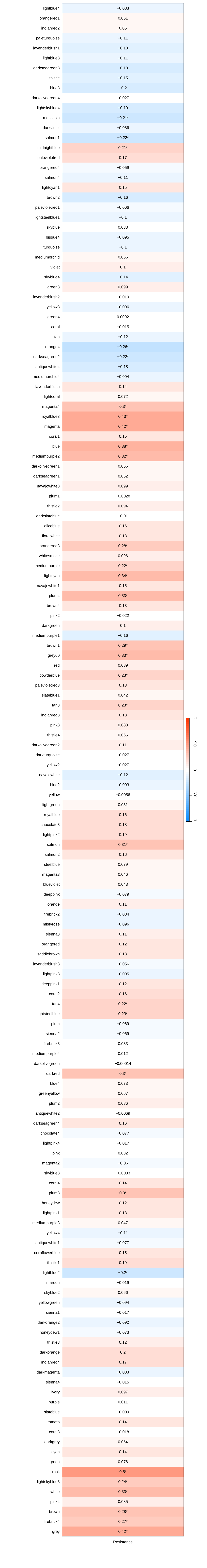
