## Supplementary File 11 for "Genome-Wide in silico analysis reveals activation of a silent resistome driving imipenem resistance in *Pseudomonas aeruginosa*"

**Table S1.** Key hub genes identified in the black module, along with their hub scores and functional annotations.

| **Gene** | **Hub Score** | **Annotation** |
| --- | --- | --- |
| group_10738 | 0.51228129 | Single-stranded DNA-binding protein; hypothetical protein |
| group_10629 | 0.51228129 | Integrase;Integrase regulator R |
| group_9158 | 0.51228129 | Receptor protein-tyrosine kinase |
| group_11099 | 0.506970866 | Cobyrinic acid ac-diamide synthase |
| group_7001 | 0.506970866 | Cobyrinic acid ac-diamide synthase;Chromosome partitioning protein |
| group_6594 | 0.506970866 | ParB/Sulfiredoxin domain-containing protein |
| group_293 | 0.506970866 | Transcriptional regulator |
| group_11177 | 0.506510038 | DUF2857 domain-containing protein;Coproporphyrinogen III oxidase |
| group_9553 | 0.474845598 | CdiI immunity protein domain-containing protein;hypothetical protein |
| group_9212 | 0.474845598 | ABC transporter substrate-binding protein;hypothetical protein |

**Table S2:** Additional hub genes identified within the respective modules containing the four genes commonly detected by both GWAS and WGCNA analyses.

| **Gene** | **Annotation** | **Module name** |
| --- | --- | --- |
| group_11241 | TraG N-terminal Proteobacteria domain-containing protein;conjugal transfer protein TraG N-terminal domain-containing protein;Conjugal transfer protein TraG;TraG-like protein N-terminal region | darkred |
| group_11116 | MobH family relaxase;Relaxase;Uncharacterized domain-containing protein;conjugal transfer nickase/helicase domain-containing protein | darkred |
| group_10871 | conjugative transfer ATPase | darkred |
| group_10870 | Conjugal transfer protein;Putative secreted protein;TIGR03751 family conjugal transfer lipoprotein | darkred |
| group_10885 | integrating conjugative element protein | darkred |
| group_10883 | Integrating conjugative element protein;TIGR03756 family integrating conjugative element protein | darkred |
| group_10882 | TIGR03757 family integrating conjugative element protein;Integrating conjugative element protein PFL_4709 family;Integrating conjugative element protein | darkred |
| group_10877 | JAB domain-containing protein;DNA repair protein RadC;MPN domain-containing protein;DNA binding protein possibly associated to lesions | darkred |
| group_3450 | hypothetical protein | plum3 |
| group_4946 | three-Cys-motif partner protein TcmP;Three-Cys-motif partner protein TcmP | plum3 |
| group_4565 | saccharopine dehydrogenase C-terminal domain-containing protein;Homospermidine synthase | plum3 |
| group_3448 | site-specific integrase | plum3 |
| group_3447 | DUF5131 domain-containing protein | plum3 |
| group_3446 | EamA family transporter | plum3 |
| group_3389 | Uncharacterized protein CL-16 | plum3 |
| group_3449 | Cointegrate resolution protein T | plum3 |
| group_4964 | Tn3 family transposase | plum3 |
| phzA | phenazine biosynthesis protein PhzA | grey |
| group_9831 | Helix-turn-helix domain protein;HTH cro/C1-type domain-containing protein | grey |
| group_12537 | Rubredoxin | grey |
| group_9685 | XRE family transcriptional regulator;HTH cro/C1-type domain-containing protein;helix-turn-helix transcriptional regulator | grey |
| group_12028 | hypothetical protein | grey |
| group_4836 | elongation factor Tu;Elongation factor Tu;EF-Tu/IF-2/RF-3 family GTPase | grey |
| group_3383 | hypothetical protein | grey |
| group_7708 | Acetyltransferase PA3944 | grey |
| rubA2 | Rubredoxin-2 | grey |
